# Fully Automated Multi-Grid Cryo-EM Screening using Smart Leginon

**DOI:** 10.1101/2022.07.23.501225

**Authors:** Anchi Cheng, Paul Kim, Huihui Kuang, Joshua H. Mendez, Eugene Y.D. Chua, Kashyap Maruthi, Hui Wei, Anjelique Sawh, Mahira F. Aragon, Viacheslav Serbynovskyi, Kasahun Neselu, Edward T. Eng, Clinton S. Potter, Bridget Carragher, Tristan Bepler, Alex J. Noble

**Affiliations:** Simons Electron Microscopy Center, New York Structural Biology Center, New York, NY, USA; Simons Machine Learning Center, New York Structural Biology Center, New York, NY, USA; Department of Biochemistry and Molecular Biophysics, Columbia University, New York, NY, USA

**Keywords:** cryo-electron microscopy (cryoEM), grid screening, automation, structural biology, computer vision, machine learning

## Abstract

Single particle cryo-electron microscopy (cryoEM) is a swiftly growing method for understanding protein structure. With increasing demand for high-throughput, high-resolution cryoEM services comes greater demand for rapid and automated cryoEM grid and sample screening. During screening, optimal grids and sample conditions are identified for subsequent high-resolution data collection. Screening is a major bottleneck for new cryoEM projects because grids must be optimized over several factors, including grid type, grid hole size, sample concentration, buffer conditions, ice thickness, and particle behaviors. Even for mature projects, multiple grids are commonly screened to select a subset for high-resolution data collection. Here, machine learning and novel, purpose-built image processing and microscope-handling algorithms are incorporated into the automated data collection software, Leginon, to provide an open-source solution for fully automated, high-throughput grid screening. This new version, broadly called Smart Leginon, emulates the actions of an operator in identifying areas on the grid to explore as potentially useful for data collection. Smart Leginon Autoscreen sequentially loads and examines grids from an automated specimen exchange system to provide completely unattended grid screening across a set of grids. Comparisons between a multi-grid Autoscreen session and conventional manual screening by five expert microscope operators are presented. On average, Autoscreen reduces operator time from ∼6 hours to <10 minutes and provides a comparable percentage of suitable images for evaluation as the best operator. Smart Leginon’s ability to target holes that are particularly difficult to identify is analyzed. Finally, Smart Leginon’s utility is illustrated with three real-world multi-grid user screening/collection sessions, demonstrating the efficiency and flexibility of the software package. Smart Leginon’s fully automated functionality significantly reduces the burden on operator screening time, improves the throughput of screening, and recovers idle microscope time, thereby improving availability of cryoEM services.

## Introduction

Over the past decade, single particle cryo-electron microscopy (cryoEM) has become an established method for structure determination of macromolecular protein complexes ranging from ∼40 kilodaltons to several megadaltons^1, 2^. A single particle cryoEM project begins with application of an aliquot of purified protein in solution to a holey foil substrate supported by a metal mesh, referred to as an EM grid. The bulk sample is then reduced to a thin aqueous film, which is vitrified by plunging the grid into a cryogen^3^. The ideal outcome of this procedure is to have the proteins spread out as ’single particles’ embedded in vitreous ice that is only slightly thicker than the largest diameter of the protein and at a concentration that enables the most efficient data collection^4, 5^. Producing suitable grids for high-resolution data collection almost always entails a series of optimization steps, with cryoEM screening required at each step to empirically examine the grids^6^. Variables that can be optimized include grid mesh type (typically copper or gold), grid film substrate (typically carbon or gold), grid hole size, sample concentration, buffer conditions, ice thickness, additives (e.g. detergents), and particle behavior like preferred orientation and degradation^3, 5, 7^. The effects of these variables on grid and sample quality requires that grids be examined in a cryo-transmission electron microscope (cryoTEM) at a series of magnifications, also called multi-scale imaging (MSI)^8^, from a grid atlas composed of grid tile images to identify squares, to Square magnification to identify regions inside squares, to Hole magnification to identify holes in those regions, to Exposure magnification to analyze protein behavior and quality (Supplemental Figure 1). MSI screening allows the operator and researcher to estimate how many images of a quality suitable for high-resolution structure determination may be obtained from each grid.

Screening across these variables usually requires that significantly more grids be prepared and imaged than are used for a subsequent high-resolution data collection, particularly for new projects. Even for mature projects, poor reproducibility of grid quality typically requires that two to six or more grids are screened before settling on a small subset that are best for a long data collection on a high-end instrument. Current data collection software packages available to the public (e.g. Leginon^8, 9^, SerialEM^10^, UCSFImage4^11^, TFS EPU^12, 13^, Gatan Latitude, JEOL JADAS^14^, AutoEMation^15^) focus on exhausting the usable imaging area on a single grid. This is commonly achieved through a high degree of tuning of automated targeting parameters. The wide range of grid types, ice thicknesses, and other confounding variables have prevented a general, robust automated solution from being developed that performs as well as an expert human operator in multi-grid screening. As a result, the major burden on a microscope operator’s time is grid screening.

To address this problem, we have incorporated a machine learning (ML) approach into our data collection system, Leginon^8, 9^, together with significant updates to Leginon’s grid handling and image processing algorithms to provide a fully automated screening application with the goal of obtaining a set of images that can be used to assess overall grid quality and identify the best regions of each grid in the microscope. The ML and some of the computer vision algorithms described herein are part of the Ptolemy package which has been described in detail elsewhere^16^, while the additional Leginon image processing algorithms are described in the Materials and Methods. We broadly call this new version Smart Leginon. ‘Smart’ refers to our effort in reducing human intervention, where the incorporation of Ptolemy square and hole targeting for automated screening is our first step. Smart Leginon includes a simple command line workflow, called Autoscreen, that allows for an entire multi-grid screening session to be set up in <10 minutes and run fully unattended. Additionally, Smart Leginon functionality may be used as independent modules from within the existing Leginon GUI. All software and algorithms described herein are free (Ptolemy is for academic use only), open-source, designed to be transferable to other collection software, and designed to be extended with new functionality.

We measured the performance of Smart Leginon in a variety of situations. First, Smart Leginon Autoscreen was used to screen 11 previously unseen mouse apoferritin (mApof)^17^ grids to assess the overall speed and robustness of the system in comparison to five expert human operators. To assess the outcomes, we measured the total screening time, total operator time, percent of good holes selected, ice thickness, and CTF resolution estimates. Next, we assessed Smart Leginon’s ability to successfully target on a wide range of grid types without adjusting any parameters, including gold and carbon substrates, multiple hole sizes and spacings, and a wide range of ice thicknesses and grid quality. Finally, we report on the application of Smart Leginon to three real-world multi-grid screening and collection sessions for users at our cryoEM facility.

## Materials and Methods

### Implementation in Leginon

In order to integrate Ptolemy into Leginon and add the Autoscreen features, three additions were built into Leginon: 1) A workflow for running Ptolemy processes which returns segmentations, target coordinates, target scores, and other metadata, 2) An algorithm to filter and sample the targets found by Ptolemy, and 3) A method to manage grid exchange and the MSI workflow for each grid. These modifications, described below, are available as of the myami-3.6 release (http://leginon.org).

Two new node classes were added to Leginon to utilize Ptolemy (1). The MosaicScoreTargetFinder handles the lowest magnification images (i.e. grid tile images of multiple squares; Supplemental Figure 1, upper-left image) which executes Ptolemy’s lowmag_cli.py on each tile of the grid atlas. Similarly, the ScoreTargetFinder node class handles Hole magnification images (i.e. images of multiple holes; Supplemental Figure 1, bottom-left image), which executes Ptolemy’s medmag_cli.py and loads the results for processing. Each of these node classes then loads the full set of results from Ptolemy into Leginon in json format. Shell scripts are defined by the Leginon administrator to make Ptolemy calls so that it is possible to easily substitute Ptolemy with future versions. An additional step was added to merge together partial squares at the edges of adjacent tiles (Supplemental Figure 2). This extra merging step allows square targets to be evaluated on the full atlas image. The merged area becomes the sum of areas, the center of gravity becomes the merged target coordinate, and the average mean intensity weighted by each target area gives the mean intensity of the merged square. The merged target also takes on the highest score of the targets it is merged from.

Atlas grid tile images are filtered by considering only squares that are within a defined square area range, while the filtering for Hole magnification images includes the Ptolemy score’s lower threshold and the ice thickness filter as implemented in other Leginon TargetFinder node classes (2). For Autoscreen purposes, Hole magnification filters are usually loosely set so only very bad selections such as holes with cracks or with large ice crystal contamination are eliminated. The Hole magnification filters did not remove any potential targets in the mApof eleven grid screening comparison.

Ptolemy scores play their strongest role for the square finder for the atlas grid tile images (2). The area-filtered squares are put into Ng groups based on the chosen parameter, where square area is typically chosen as in the results presented herein. The highest-ranked (‘best’) squares from each group based on Ptolemy scores then creates a total of Ns squares to be targeted at higher magnifications. For example, if Ng = 4 and Ns = 8, then 2 squares from each group will be selected.

A sampling feature for the Hole magnification image hole finder was added to Leginon’s automated target finder base class (2). This presents the user with one additional setting to handle the sampling and to decide on the maximum number of targets Nh to include. Sampling in a given Hole magnification image is produced by dividing the holes by a defined variable into Nh classes and then randomly sampling one instance in each class. Relative ice thickness as determined by average pixel intensity of a small group of pixels near the center of a hole is used as the variable in this classification. The Ptolemy scores can also be used to filter the targets prior to sampling, however this filter was not used in any of the results presented herein.

The Autoscreen workflow is initiated by a command line python script that sets up session information and defines the task to perform on each session (3). The current options are “full MSI”, which performs unattended grid screening at all magnifications, and “atlas only”, which only collects an atlas for each grid. This required additional changes in the Leginon framework. Changes to the Leginon manager were made to switch sessions without disconnecting from instruments and to issue the grid exchange and workflow instruction to individual sessions when it is active. Settings were made to be recallable from a specified example session instead of the most recent session. Automated execution of the square finder was added as an option for the MosaicTargetFinder base class. A “Center between holes” option was added to the current auto-creation of focus targets. This parameter-less algorithm analyzes the target lattice and places the focus target halfway between the lattice points nearest to the center of the Hole magnification image, thus ensuring its maximal distance from any hole selections.

To fully realize unattended multi-grid screening with Autoscreen, we use AutoIt scripting (https://www.autoitscript.com) to emulate the GUI operations necessary to insert and retract the objective aperture in the TFS microscope API as these function calls are not available through the TFS microscope API (3). For the examples shown in this manuscript, Ptolemy was run on a single CPU core on the Leginon computer connected to the microscope, sufficient to keep up with real-time collection.

### Smart Leginon Autoscreen workflow

Figure 1 illustrates the general Smart Leginon workflow, with Autoscreen functionality highlighted in blue. The operator provides Autoscreen with a list of grids to be screened in the order in which they should be imaged and associates each grid with a specific project in the database. The session is started after which the following actions are performed unattended: (i) a grid is loaded from the automated specimen exchange system into the microscope, (ii) an atlas of the entire grid is collected tile-by-tile, (iii) Ptolemy locates all squares in the atlas tile images and merges the results, (iv) Leginon separates squares into Ng groups based on a chosen parameter (currently either square area, mean intensity, or Ptolemy score), and chooses the highest-ranked square from each group for a total of Ns squares, (v) Leginon acquires a Square magnification image of each targeted square, then a raster of Hole magnification images are acquired after the grid is set to eucentric height in the square, (vi) Ptolemy identifies the positions of holes within each Hole magnification image, (vii) Leginon selects Nh holes from this set based on chosen parameters (e.g. ice thickness, Ptolemy score, and/or random), (viii) Leginon performs a set of procedures to acquire a Exposure magnification image from each hole; including focusing, setting defocus, checking for drift, and normalizing lenses, (ix) Once all squares and all holes from a grid have been imaged, the grid is unloaded from the stage. This process is then repeated until all grids have been examined, after which Autoscreen is terminated and a message is sent to the operators and users, typically through a designated Slack channel (https://slack.com). During grid exchange, Leginon retracts and inserts the objective aperture automatically. Leginon performs ice thickness estimation^8, 18^ for each Exposure magnification image in real-time. After each grid is screened, Smart Leginon initiates frame alignment and CTF estimation through Appion^19^. These image processing procedures are fast enough to be performed at the same rate as data collection. Exposure magnification collection targets may be augmented by collecting at several locations away from or in addition to the center of the holes identified by Ptolemy; i.e. multi-shot hole targeting. All of the images and the pre-processing results can be viewed in the Appion^19^ web-based 3 Way Viewer (Supplemental Figures 1 & 3). Several components of the Autoscreen functionality may be used as independent modules from within the Leginon GUI, e.g. most hole finding at SEMC is now done using the Ptolemy hole finder.

**Figure 1:**
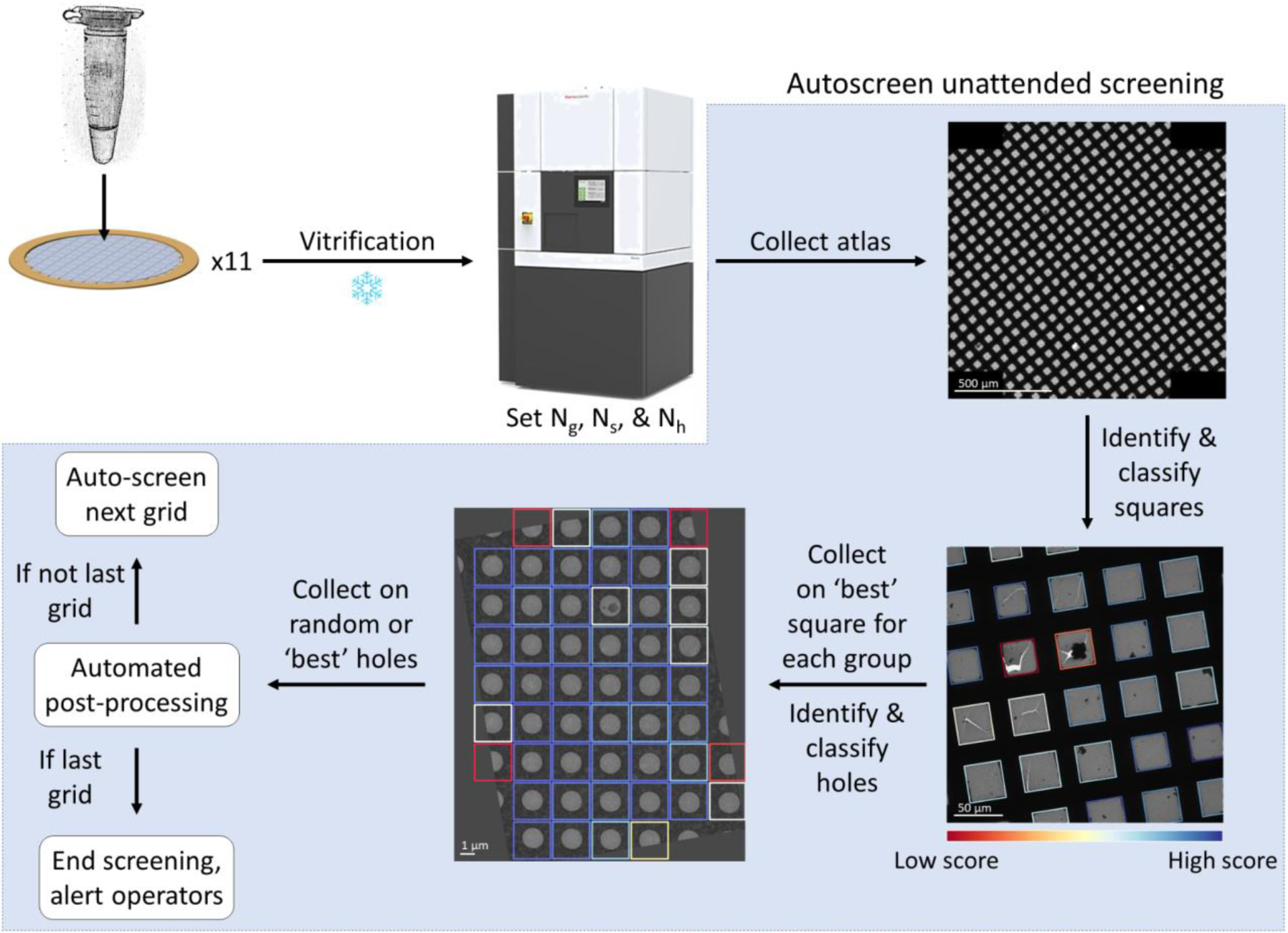
Smart Leginon Autoscreen fully automated multi-grid screening workflow. The general Smart Leginon Autoscreen fully automated, unattended multi-grid screening workflow. CryoEM grids with samples are prepared and loaded into a cryoTEM with an automated specimen exchange system. In Autoscreen, the microscope operator imports settings from an example session, including 1) Ng, the number of groups to separate the squares in the grid atlas by, based on square area, mean intensity, or Ptolemy score, 2) Ns, the total number of squares that will be collected, and 3) Nh, the number of holes that will be collected for each square. For each grid in the microscope, Smart Leginon Autoscreen will automatically target the highest-ranked (‘best’) square in each group and sample Nh random, threshold-filtered holes for each square, where Ptolemy determines rankings. Exposure magnification ice thickness estimations are determined in real-time by Leginon and automated pre-processing in Appion (e.g. frame alignment and CTF estimation) is initiated at the end of each grid collection. If another grid is listed for screening, then Autoscreen automatically moves to the next grid and initiates a new collection session in the database. Once all grids are screened, Autoscreen safely returns the microscope to its default state and sends a message of completion to a designated Slack channel to alert operators and users. Each step in Autoscreen may be performed unattended by command line or used as independent modules in the Leginon GUI. Atlas scale bar is 500 μm, grid tile image scale bar is 50 μm, hole image scale bar is 1 μm.

### Mouse apoferritin cryoEM grid preparation and screening

Eleven cryoEM grids were prepared by two people by adding 3 μL of mouse apoferritin (mApof, from Dr. Kikkawa’s lab) solution (8 mg/mL) to UltrAuFoil 1.2/1.3 grids (Quantifoil, Jena, Germany) immediately after plasma cleaning (Gatan Solarus II plasma cleaner; Gatan Inc., Pleasanton, CA, USA). The grids were blotted for 4 or 4.5 seconds, then vitrified by plunge-freezing in liquid ethane using a TFS Vitrobot Mark IV (Thermo Fisher Scientific) with the chamber maintained at 20 °C and 100% humidity.

The eleven mApof cryoEM grids were screened on a TFS Glacios with a Falcon 3 camera (Thermo Fisher Scientific) in integration mode. The grids were not pre-screened prior to loading into the Glacios and starting Smart Leginon. Parameters used were Ng = 4, Ns = 4, and Nh = 5 where these and all other settings were imported from a previous screening session that created an example Smart Leginon session (Supplemental Figure 4). The atlases consisted of 22 grid tile images each, where each tile was acquired at a magnification of 210x (2751 Å/pix). Square magnification was set to 940x (615 Å/pix); Hole magnification was set to 5,300x (109 Å/pix); Exposure magnification was set to 120,000x (1.204 Å/pix). Exposure magnification movies were recorded in linear mode with a total exposure time of 400 ms across 40 frames and with an accumulated electron dose of 55.59 e^-^/Å^2^ at -3 μm nominal defocus.

### Smart Leginon Autoscreen versus operators quantification metrics

Timing measurements for Smart Leginon Autoscreen and human operators were obtained by using the image timestamps at the beginning and end of each session. To reduce the bias of external microscope hardware, the time required for microscope alignment before the screening session and LN2 fillings were removed from time measurements. Grid exchange times (∼5 minutes to retract a grid and insert the next grid) are included in the time measurements.

Hole quality analysis for mApof grids was performed visually. Only holes with greater than ∼80% of the hole area existing inside the image (i.e. not significantly cut off by the edge of the image) were considered. A hole was considered contaminated if greater than ∼40% of the imageable area in the hole was obfuscated.

### Real-world Smart Leginon Autoscreen 35-grid user session

An assortment of grids - Quantifoil R1.2/1.3 300 mesh, Quantifoil R1.2/1.3 300 mesh with graphene, UltrAuFoil R1.2/1.3 300 mesh (Quantifoil, Jena, Germany) - were frozen with a TFS Vitrobot Mark IV (Thermo Fisher Scientific). 35 user grids were screened on a TFS Glacios with a Falcon 3 camera in integration mode. Autoscreen settings were Ng = 3, Ns = 3, and Nh = 3. Imaging parameters were the same as the mApof screening session, except the atlases consisted of 43 grid tile images each and Square magnification was set to 2,600x (222 Å/pix).

### Real-world Smart Leginon user sample screening and collection session

Eight user grids were screened on a TFS Glacios with a Falcon 3 camera in integration mode. Autoscreen settings were Ng = 4, Ns = 4, and Nh = 5. Imaging parameters were the same as the mApof screening session, except the atlases consisted of 28 grid tile images each. The two best grids were selected for a full data collection on a TFS Krios with a Gatan K3 camera in counting mode and BioQuantum energy filter (Gatan Inc., Pleasanton, CA, USA). Hole magnification was set to 3,600x (97 Å/pix). Exposure magnification was set to 81,000x (1.069 Å/pix). 2D classification and 3D refinement were performed with CryoSparc v3.3.1^20^.

### Real-world Smart Leginon user data collection session

Grids were screened and collected on a TFS Krios with a Gatan K3 camera in counting mode and BioQuantum energy filter. Hole magnification was set to 3,600x (76 Å/pix). Exposure magnification was set to 81,000x (0.846 Å/pix).

### Micrograph pre-processing

Motion correction was performed with MotionCor2^21^ and CTF parameters of motion-corrected micrographs were estimated by CTFFIND4^22^ through the Appion^19^ pipeline. Ice thickness was determined by the ALS method for Glacios sessions and by the energy filter method^18^ for Krios sessions from within Leginon^8^; estimation by ALS is accurate to an estimated ±10 nm. For Smart Leginon Autoscreen sessions, AutoRelauncher.py was used to automatically re-launch the Appion real-time pre-processing (i.e. frame alignment and CTF estimation) from an example session for many screening sessions described herein.

## Results

### Smart Leginon Autoscreen significantly decreases operator time while increasing microscope throughput

The Smart Leginon Autoscreen multi-grid screening performance using a TFS Glacios was evaluated by comparing it to five microscope operators who had not previously seen the grids nor the Autoscreen results. To obtain metrics on a per-grid basis, 11 grids were screened by Autoscreen in one Smart Leginon session to establish timing and performance values. Three of these grids were selected for evaluation by five expert operators for a total of 15 grid screenings by operators. Autoscreen and each operator targeted Ns = 4 squares across Ng = 4 groups and selected Nh = 5 random holes per square. The operators selected squares manually but set up the standard Leginon template matching hole finder to target five holes closest to the center of the image used for hole targeting. A variety of metrics were measured to assess the outcomes, including: total screening time (i.e. the time from inserting the first grid into the microscope to the time the last grid is retracted), total operator time required during screening, the percent of “good” holes selected (i.e. non-empty, minimal contamination, no cracks), ice thickness estimation^18^ as measured from the Exposure magnification images, and CTF resolution estimation^23^.

The Autoscreen collection sessions, including the example Smart Leginon session, took about 10 minutes for the operator to set up before beginning unattended collection. Screening of each grid then took an average of 29.7 ± 1.2 minutes to collect, resulting in about 5.4 hours total to screen 11 grids. The equivalent tasks performed by five expert microscope operators on three of the 11 grids took 32.7 ± 7.1 minutes per grid, which extrapolates to 6.0 hours to screen 11 grids if the operator rarely or never leaves the microscope. While screening grids by an operator generally requires the operator to stay at the microscope during the entire process in order to perform several manual tasks, there does exist some time for the operator to non-optimally multitask. The amount of time depends strongly on the quality of the grids, which is unpredictable. During these tests, each operator was not interacting with the microscope for 5- 10 minutes of fragmented time per grid, which is generally not enough time to accomplish any meaningfully-involved task. Thus, we estimate that the microscope operators have little to no meaningful time away from the microscope during their entire screening session. Figure 2 summarizes the results.

**Figure 2:**
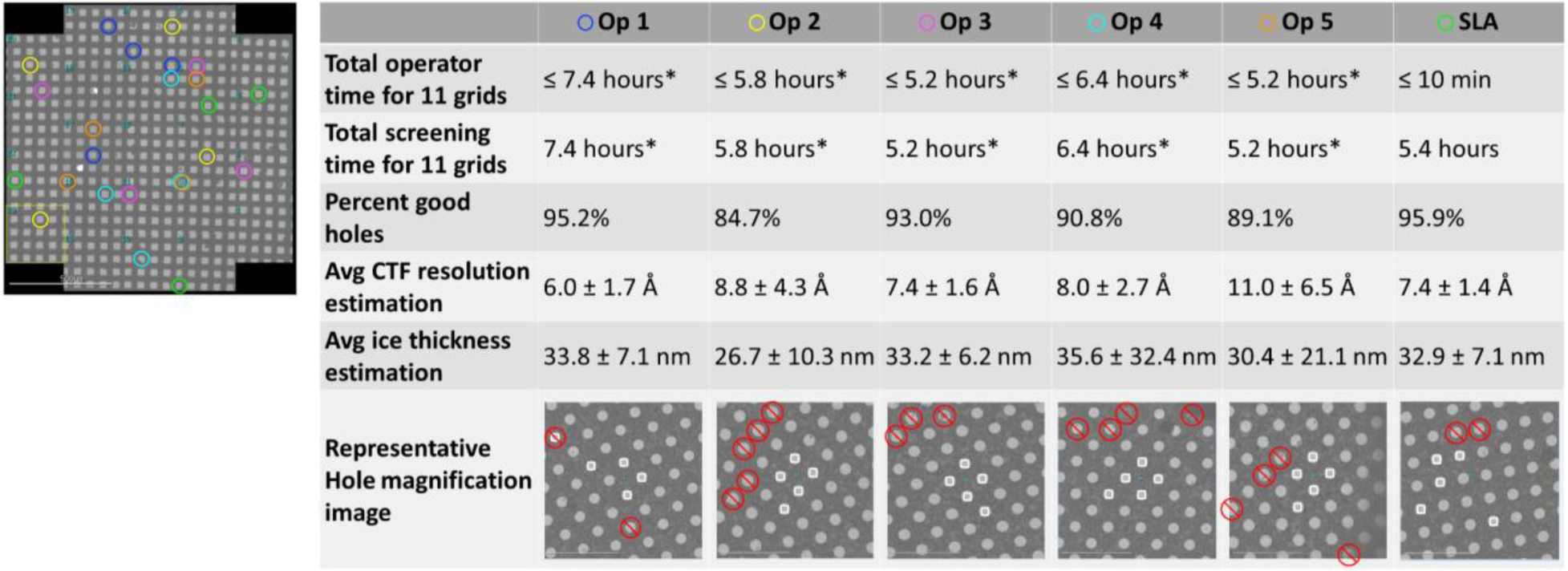
Smart Leginon Autoscreen comparison against expert microscope operators. Quantitative comparisons between Smart Leginon Autoscreen (SLA) and expert microscope operators (Op). Autoscreen and operators each independently selected what they considered to be the ‘best’ 4 squares, where for Autoscreen each square was selected from four equally-spaced square area ranges spanning all square areas (one grid atlas is shown on the left; see Supplemental Figure 8 for all atlases). Autoscreen took <10 minutes for the operator to set up and then run in a completely unattended manner for 5.4 hours. The operators spend an average of 6.0 hours to screen 11 grids (*extrapolated from 3 grids), of which most of the time is spent operating the microscope, interspersed with several short periods (5-10 minutes per grid) of time away from the microscope. (Note: Calculations assume that the operators do not take any breaks away from the microscope.) In terms of percent of good holes available from Hole magnification images, Autoscreen (95.9% good holes) performed better than the average from the operators (90.6% good holes) and comparable to the best operator (95.2% good holes). Supplemental Figures 9-14 show visual analyses of all holes and Table 1 shows the quantifications. From the random holes targeted, CTF resolution estimation for Autoscreen holes (7.3 ± 2.6 Å) was comparable to the average obtained by operators (7.4 ± 2.9 Å). Estimates of ice thickness show comparable values between Autoscreen (32.9 ± 7.1 nm) and operators (35.0 ± 18.7 nm). Supplemental Table 1 shows the raw data. The Hole magnification image most representative of the average in terms of hole quality is shown in the last row. Targets are shown by white squares and bad holes are shown by red circles with lines through them. Note: The representative image for Operator 4 is a composite image due to there not existing an image that closely represents the average.

The Autoscreen images at all magnifications were visually inspected in the web-based viewer that is part of the Leginon and Appion package^19^ (Supplemental Figure 1) which quickly allowed the operator to determine that eight of the grids were of good to excellent quality while the other three were of poor quality (Supplemental Figure 5). This analysis took about five minutes and allowed the operator to correctly correlate which grids were made by which person, exemplifying the efficiency of combining automated screening software with a database for storing data and a web GUI for rapid, remote visual analysis.

### Smart Leginon square grouping and ranking replicates operator performance

One goal of cryoEM grid screening is identifying squares where optimal holes reside. For *de novo* cryoEM projects where no cryoEM screening has been performed, a common practice is to screen several different squares with different visible areas because the square area is often inversely proportional to the ice thickness of holes in the square (Supplemental Figure 6). On the other hand, if the sample owner has prior knowledge of the optimal square area for their sample, then the microscope operator will concentrate screening efforts on those squares.

Smart Leginon can be optimized for different stages of a project. Large values of square groups, Ng, ensures that diversity is achieved in *de novo* cryoEM projects in a manner comparable to microscope operators. If the sample owner has prior knowledge of the sample’s behavior in ice, then the square area can be restricted to a range and Ng can be set to ‘1’ so that only the highest-ranked Ns squares within a specific area range are collected (Supplemental Figure 7).

Supplemental Figure 8 shows the three grids that were screened by Smart Leginon Autoscreen and independently by the five expert microscope operators. There is nearly no overlap (2.8% overlap) between the squares identified by Autoscreen and the operators, which is likely due to the fact that there are a large number of possible squares in each group area range and that several squares in each group are visually indistinguishable. Manual examination of each Hole magnification image for Autoscreen and operator collections was performed to identify “bad” holes (i.e. holes that have no ice, that have considerable contamination, or that have cracks) (Table 1, Supplemental Figures 9-14). Smart Leginon Autoscreen using Ptolemy performed better at finding squares with good holes (95.9% good holes) compared to average operator performance (90.6% good holes).

**Table 1:**
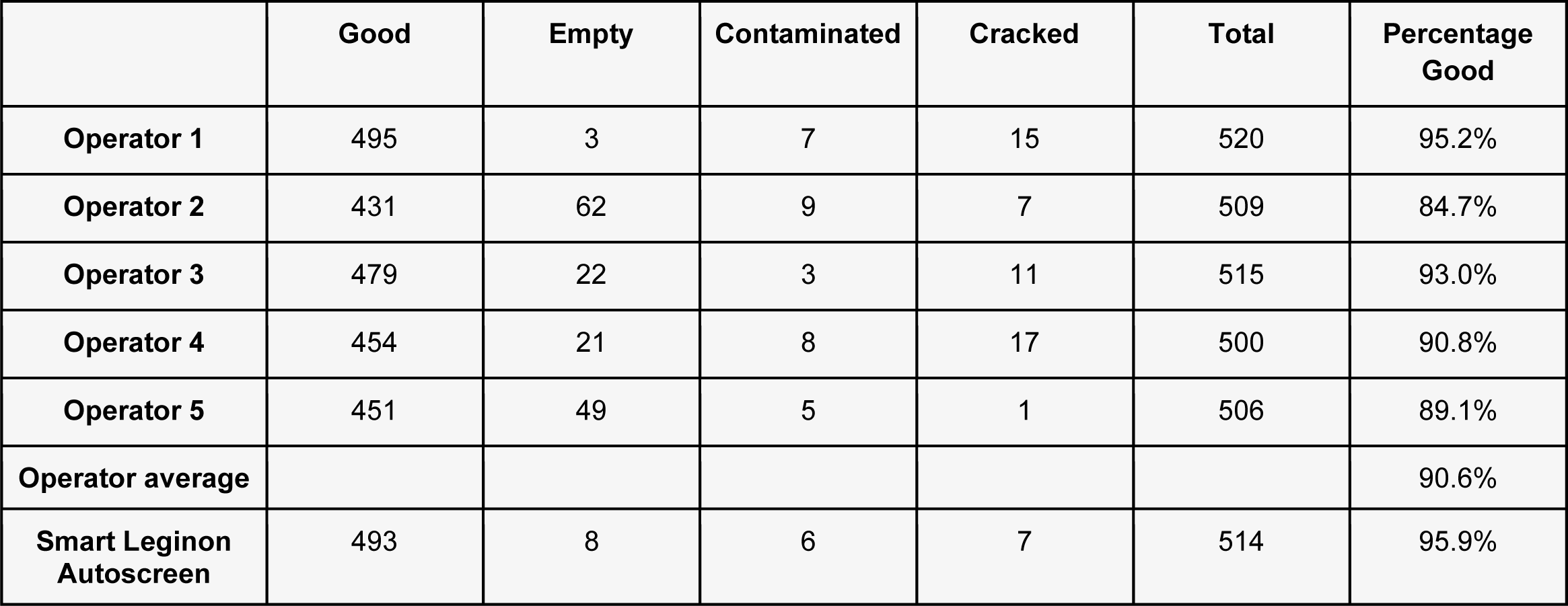
Analysis of holes from each square image. Hole analysis for each square image for the three mApof grids that were imaged by Smart Leginon Autoscreen and the five expert microscope operators. Supplemental Figures 9-14 show the annotated Hole magnification images.

This same level of performance was also evident when the Exposure magnification images were analyzed. CTF resolution estimation of all Exposure magnification images showed comparable performance between Smart Leginon Autoscreen (7.3 ± 2.6 Å) and operators (7.4 ± 2.9 Å). Ice thickness estimates for Smart Leginon Autoscreen (32.9 ± 7.1 nm) showed a comparable average and narrower range compared to the operators (35.0 ± 18.7 nm) (Figure 2, Supplemental Table 1).

### Smart Leginon identifies holes independent of grid type and hole size

Smart Leginon hole lattice identification performance using Ptolemy was tested on multiple different types of grids. We found that Ptolemy hole finding for carbon film (Figure 3a-d) and gold film (Figure 3e-h) grids with varying hole sizes and spacings generally performs well without the need to adjust any parameters. Ptolemy hole targeting performed well under conditions where the template matching hole finder would have struggled or failed, for example on images where holes are darker than the surrounding film (Figure 3b), on thick ice images with low contrast between the holes and foil (Figure 3c), and on images with contamination (Figure 3a-d).

**Figure 3:**
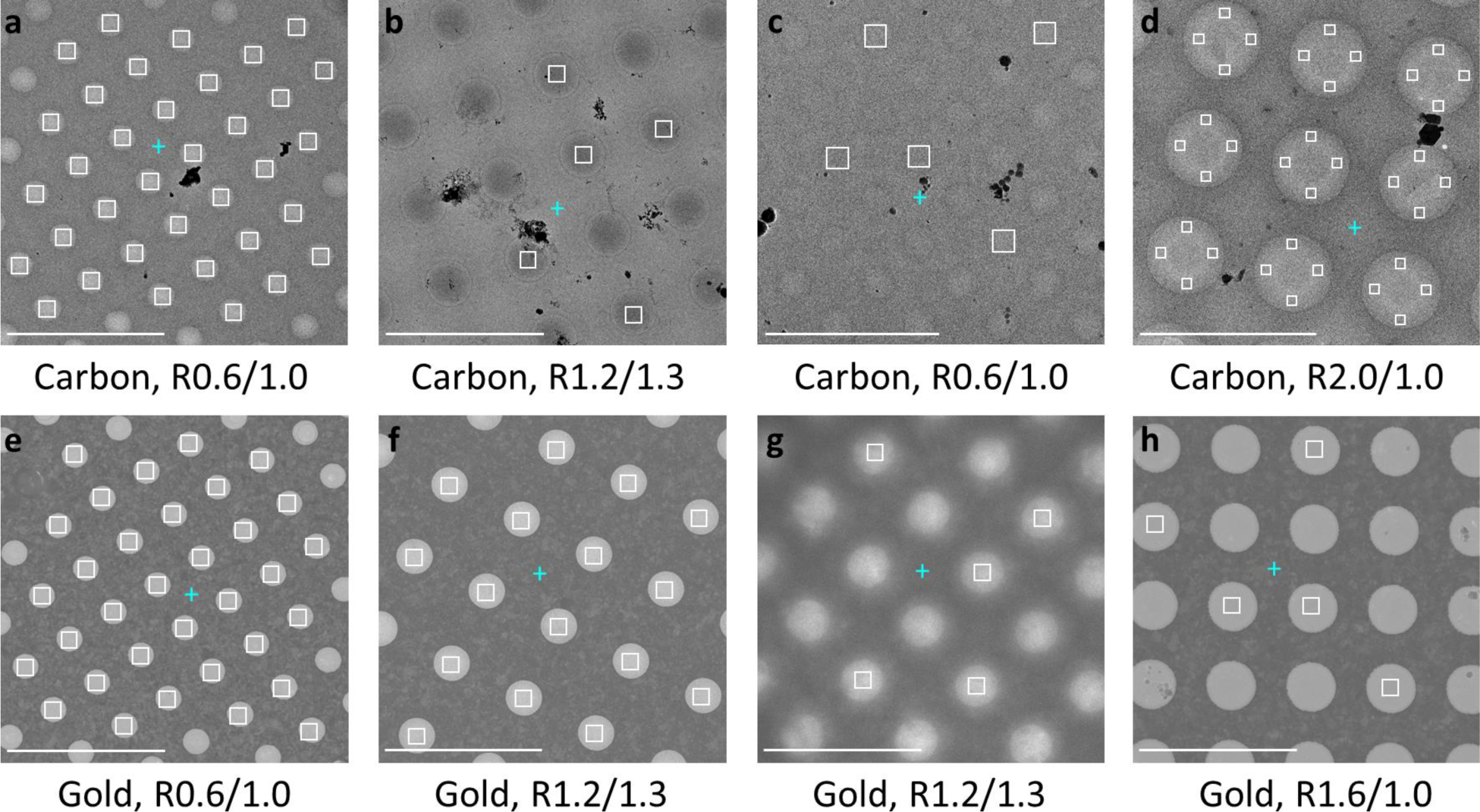
Smart Leginon hole targeting applied to different grid types. Smart Leginon hole finding using Ptolemy applied to Quantifoil grids of different material, hole size and spacing, and characteristics. The top row shows targeting on carbon foil grids and the bottom row on gold foil grids. **(a)** An R0.6/1.0 carbon grid with minor contamination and uniform ice where all holes were identified and targeted (white field of view boxes). **(b)** An R1.2/1.3 carbon grid with moderate contamination where the holes are darker than the surrounding foil and where all holes were identified and a random subset of five targeted. **(c)** An R0.6/1.0 carbon grid with thick and non-uniform ice and minor contamination where all holes were identified and a random subset of five targeted. **(d)** An R2.0/1.0 carbon grid with minor contamination where all holes were identified and targeted with four multi-shot targets. **(e)** An R0.6/1.0 gold grid with minor contamination (top-left hole) where all holes were identified and all non-contaminated holes targeted. **(f)** An R1.2/1.3 gold grid with no contamination where all holes were identified and targeted. **(g)** An R1.2/1.3 gold grid with thick ice and no contamination where all holes were identified and a random subset of five targeted. **(h)** An R1.6/1.0 gold grid with minor contamination where all holes were identified and a random subset of five targeted. Blue plus signs (+) are automatically-determined focus locations. Note: An exclusion border was applied to each image, so some edge holes are not targeted. Scale bars are 5 μm.

### Smart Leginon and Autoscreen applied to a real-world, 35-grid screening session

Smart Leginon Autoscreen was used to automatically screen 35 grids across three samples from one user on a TFS Glacios over four intensive days of concurrent grid optimization. Various grid types (carbon substrate, gold substrate, and graphene-coated grids), sample concentrations, and grid-making conditions (i.e. changes in blotting time on the Vitrobot) were attempted for each sample with the goal of preparing and identifying Krios-ready grids. Generally, grids were prepared during the day and screened automatically with Autoscreen, usually overnight, then the screening data was evaluated the next morning and used to guide the next iteration of grid and sample preparation. To prepare for screening all grids, the first grid was screened semi-manually while determining suitable Smart Leginon parameters to create an example session for Autoscreen. On two occasions, Autoscreen completed before the end of the working day, which allowed for preliminary data collection on the Glacios to be collected on the best grid overnight using Smart Leginon and the Ptolemy hole finder. 2,594 micrographs were collected during one of these unattended overnight sessions resulting in a 5.6 Å structure, allowing for verification of the quality of the grid-making conditions for this sample. With an average automated screening time of ∼29 minutes per grid, Autoscreen and Smart Leginon enabled 35 grids to be automatically (for 34 grids) and semi-automatically (for the example grid) screened over ∼17 hours of microscope time (overnight collection not included) during the 4-day period, allowing for a constant rapid feedback loop to the grid-making process. After all grids were prepared and screened, 12 grids were determined to be ready for Krios data collection. The Smart Leginon workflows allowed for the grid preparation and screening cycle to be significantly condensed, substantially increasing the efficiency of the microscope’s, operator’s, and researcher’s time.

### Smart Leginon applied to a real-world user sample screening and collection session

Smart Leginon Autoscreen was used to *de novo* screen eight user grids of an unspecified sample on a TFS Glacios. In general, the eight grids had thick ice, with three grids being completely opaque, three grids having a very limited number of good squares, and the remaining two grids having reasonable, though thick (100+ nm), ice and a sufficient number of good squares for high-resolution collection (Figure 4a,b). The best of these two grids was transferred to a TFS Krios for a full collection. During high-resolution data collection, the user chose to manually target squares, while Smart Leginon’s implementation of Ptolemy was used for hole targeting. The grid had a wide range of ice thicknesses: particularly thick ice resulted in some holes appearing lighter than the surrounding film (as commonly seen in, e.g., Figure 3a), some holes appearing as nearly the same contrast as the surrounding film (Figure 4c), and some holes appearing darker than the surrounding film (Figure 3b). Ptolemy from within Smart Leginon allowed for holes in all cases to be identified reliably in an unattended manner without changing any parameters. The Krios session resulted in nearly 4,000 Exposure magnification images (Figure 4d) whose ice thicknesses were primarily over 100 nm (Figure 4e), yet reported good CTF resolution estimations (Figure 4f), likely due to the highly concentrated proteins (Figure 4d). Subsequent 2D classification (Figure 4g) and 3D refinement resulted in a 3.1 Å EM map (not shown here), which was sufficient for biological interpretation.

**Figure 4:**
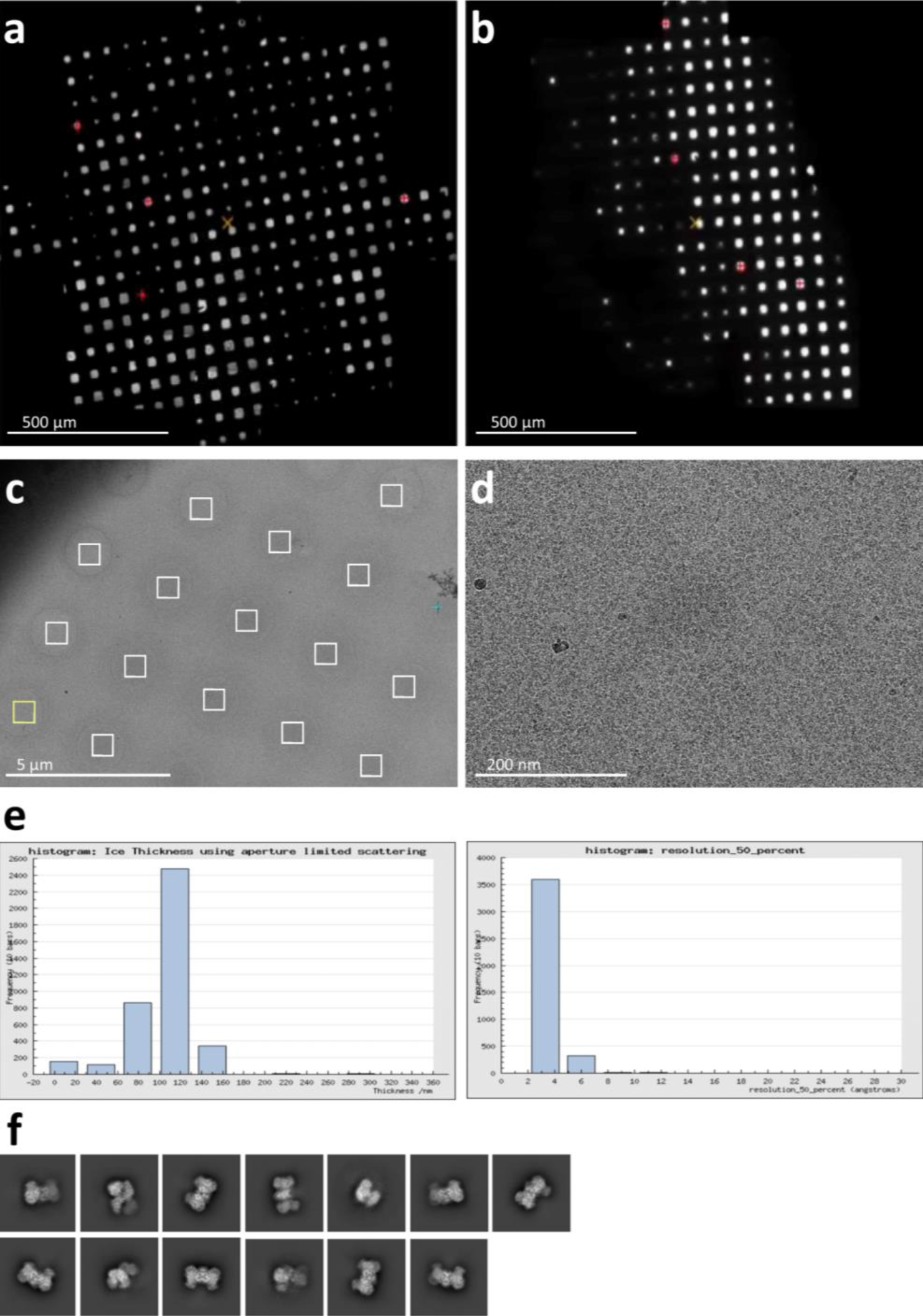
Real-world Smart Leginon multi-grid screening and collection session. Real-world multi-grid screening with Smart Leginon Autoscreen followed by data collection with the Smart Leginon and the Ptolemy hole finder. Autoscreen readily allowed for the identification of the best grid **(a)** and another decent grid **(b)**. Ptolemy’s hole finder from inside Smart Leginon was then used for high-quality data collection from the best grid **(c)**, leading to thousands of Exposure magnification images **(d)** in areas of moderately thick ice resulting in relatively high-resolution CTF estimations **(e)**, high-quality 2D classes **(f)**, and a 3.1 Å structure (not shown).

### Smart Leginon screening applied to a real-world user data collection session

A common user practice is to first screen sample and grid conditions on a screening microscope, then after variables are found that produce high-quality cryoEM grids, create several grids under the same conditions and load them directly into a high-end cryoTEM to avoid potential contamination during transfer from the screening microscope. However, the reproducibility of cryoEM grids under identical grid-making conditions is low. As a result, several grids often need to be screened in the high-end cryoTEM before ranking the grids for collection. This step again entails potentially hours of additional screening work by the microscope operator.

We employed Smart Leginon to screen four freshly-made cryoEM grids on a TFS Krios prior to high-resolution data collection. The user had ordered the grids for data collection based on blot time according to previous screening results (Supplemental Figure 15). For each grid, Ptolemy square targeting together with Smart Leginon algorithms were used to select 5 squares across a range of square areas that were thought would be most likely to contain well-behaved particles. The operator then used Smart Leginon and the Ptolemy hole finder to automatically screen Nh = 4 holes per square in an unattended manner. After ∼30 minutes of screening for each grid, the operator and user deduced that 1) the optimal areas for collection were in squares with moderate ice thickness (30-40 nm; Figure 5d; red circles in Supplemental Figure 15) rather than thin ice areas (10-20 nm; Figure 5a-c) as anticipated, and 2) the anticipated grid ordering by the user was exactly the opposite of the optimal ordering as determined by Smart Leginon MSI analysis (Supplemental Figure 15). With this knowledge, the operator queued the best 2 grids using the Ptolemy hole finder in Smart Leginon for a 43-hour collection session resulting in over 12,000 micrographs and a 2.6 Å EM map (not shown here). This use of Smart Leginon for 2 hours of screening just prior to a high-resolution data collection proved critical for maximizing the efficiency of Krios time.

**Figure 5:**
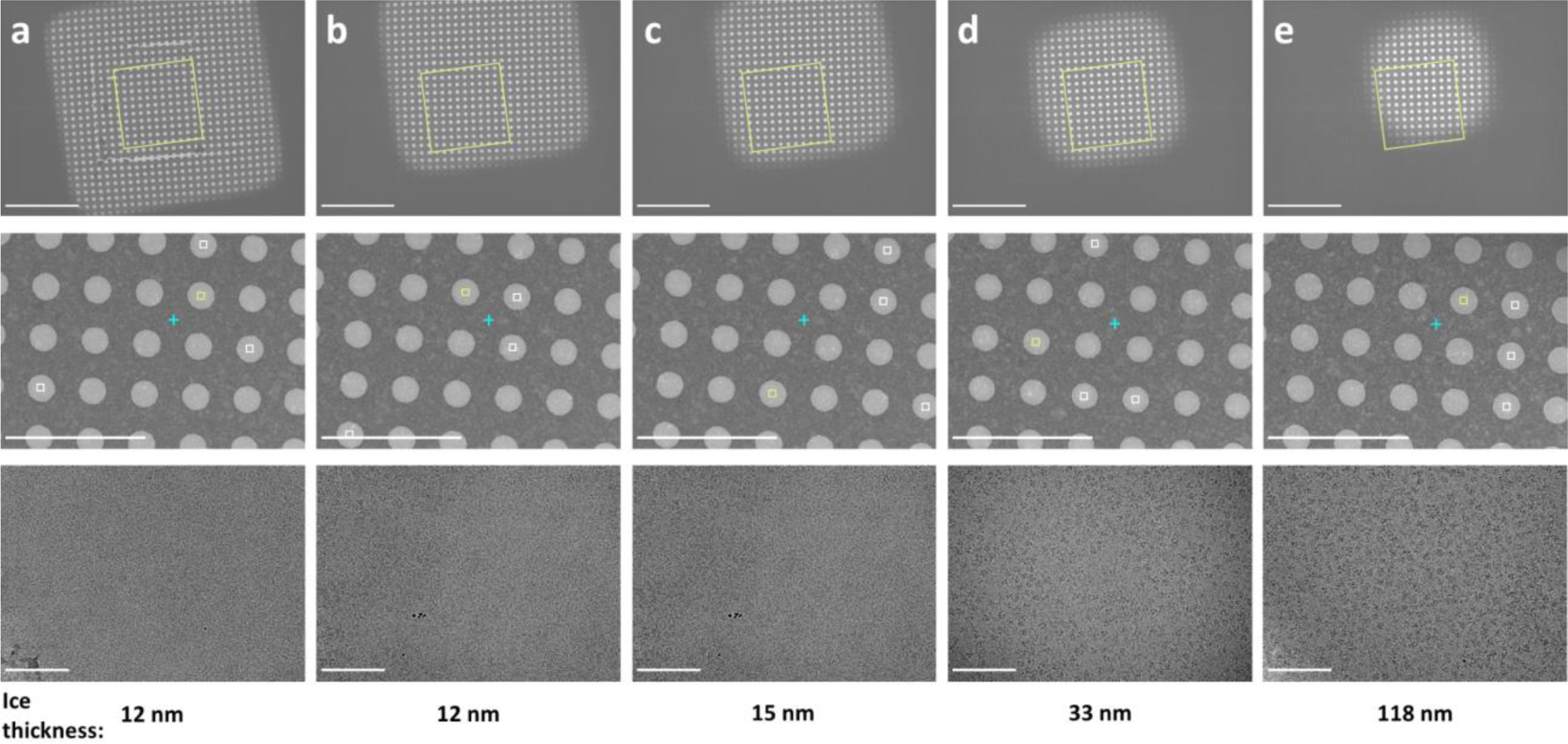
Real-world Smart Leginon multi-grid screening and collection session. Real-world Smart Leginon screening MSI grid analysis of a user’s grid. The MSI analysis, with Square magnification images of several squares from Grid 3 in Supplemental Figure 15 in the first row, Hole magnification images of corresponding holes in the second row, Exposure magnification images from inside corresponding holes in the third row, and ice thickness estimations in the fourth row, allowed the operator and user to determine that medium-sized squares contain 30-40 nm thick ice that contains non-overlapping particle of interest **(d)**, while squares with ice thinner than 30 nm ice were void of particles **(a-c)** and squares with ice on the order of 100 nm thick had overlapping particles **(e)**. Scale bars for the first row are 10 μm, 5 μm for the second row, and 100 nm for the third row.

## Discussion

Single particle cryoEM throughput is tracking x-ray crystallography throughput from 20 years ago^24^ and is poised for a throughput revolution^12^ much like the previous resolution revolution^25^. To keep up with the increasing demand from structural biologists, cryoEM developers must significantly reduce bottlenecks in the workflow. One significant bottleneck is the screening of cryoEM grids and samples prior to high-resolution collection. This step consumes a significant amount of user and microscope operator time that could be better used in other bottleneck areas such as sample preparation and data analysis. Fortunately, ML algorithms and image analysis have progressed to the point where most cryoEM screening tasks at the microscope can be performed without user intervention.

To directly address the cryoEM screening problem, we present Smart Leginon for fully automated grid screening. We illustrated the improvements of using Smart Leginon Autoscreen by experimentally testing the software against five expert microscope operators for three mApof grids and found that Smart Leginon required significantly less operator time at the microscope (<10 minutes for 11 grids compared to 6 hours) while targeting comparable squares. We also showed that Ptolemy’s parameter-less hole finder performed well on a range of common and difficult hole identification tasks with carbon and gold holey grids, including when contrast variation between the holes and the surrounding film reverses. The parameter-less hole finder suitably addresses the long-standing bottleneck of cryoEM hole identification. Lastly, we deployed Smart Leginon on three real-world multi-grid user samples to illustrate its utility; 1) A 4-day session where the user and operator iteratively screened 35 grids with Autoscreen and Smart Leginon while looping this rapid feedback into sample and grid optimization, allowing for 12 Krios-ready grids to be made; 2) A *de novo* cryoEM Autoscreen session of eight grids where it was determined that relatively thick ice contained well-behaved particles at reasonable concentration. The two best grids were collected using Smart Leginon and Ptolemy, resulting in a 3.1 Å EM map; and 3) A Krios session where four unseen grids were screened with Smart Leginon and Ptolemy, where the grid conditions were previously known and the grid order was prioritized by user based on identical grid preparation conditions. Smart Leginon Autoscreen results suggested better priority and the best two grids resulted in a 2.6 Å EM map. Additionally, Smart Leginon Autoscreen allowed for microscope idle time, including overnight and on weekends, to be used for screening due to the minimal operator time required, further increasing the overall cryoEM throughput from sample preparation to high-resolution data collection.

Smart Leginon’s improvements, particularly the reduction of operator time from 6 hours to <10 minutes for screening 11 grids, can be attributed equally to the ML algorithms and image processing in Ptolemy and the new purpose-built algorithms in Leginon. The targeting tasks have historically been performed by operators either manually or in a semi-automated manner where the operator either selects targets by hand or adjusts several parameters until the software reliably targets a narrow range of squares/holes. These specialized parameters, however, often do not translate well to new grids. Additionally, these semi-automated algorithms take time and expertise to use, and cannot account for wide variations in grid conditions, such as the grid in Figure 3b & 4c which has alternating contrast between the holes and the surrounding film. In contrast, Smart Leginon’s implementation of Ptolemy for hole finding has no parameters to change while square finding has only four parameters that need to be set at the outset of a multi-grid Autoscreen session: Ng, Ns, Nh, and the range of either square area, mean intensity, or Ptolemy score to search for squares within. Equally important, Smart Leginon Autoscreen’s ability to insert and retract grids and the objective aperture were critical for fully automating screening.

The Smart Leginon workflow (Figure 1) may be extended in multiple ways, for instance: 1) The results from the Smart Leginon workflow may be augmented by numerous live-processing software packages^20, 26–32^ that perform, for example, particle picking, ab-initio model generation, 2D/3D classification, and 3D refinement as automated post-processing routines, and 2) Targeting may be further improved by feeding live-processing results back into the collection software as targeting priors.

### Potential for implementation in other collection software

The image processing and ML routines from Ptolemy that were integrated together with the algorithms added into Leginon may also be integrated into other collection software packages, such as SerialEM^10^, UCSFImage4^11^, TFS EPU^12, 13^, Gatan Latitude, JEOL JADAS^14^, and AutoEMation^15^. We describe in the Materials and Methods the exact modifications required to integrate Ptolemy into Leginon and to perform filtering and grid manipulation. The generalized requirements are: 1) A workflow for running Ptolemy processes and returning segmentations, target coordinates, target scores, and other metadata that Ptolemy provides, 2) An algorithm to sample the targets found through automation, and 3) A method to manage grid exchange and the MSI workflow for each grid.

Once these algorithmic requirements are met, then Ptolemy can be run in real-time using a CPU on any modern computer with access to the same filesystem as the collection software. In addition to Ptolemy integration, several collection software modifications described in the Materials and Methods may need to be made, such as automatically controlling the objective aperture or integrating image processing routines for estimating clustered pixel intensities.

### Using other ML hole and square target classifiers in Smart Leginon Autoscreen

There are a few hole and square target classifiers available^33, 34^. These can be used in Smart Leginon Autoscreen as long as target finding results are provided by the classifier in the format accepted by the two ScoreTargetFinder node classes described here. For API details, please refer to https://emg.nysbc.org/redmine/projects/leginon/wiki/MSI-Ptolemy_API_information_for_developers

### Current limitations of square selection

Due to the wide and sometimes unpredictable range of grid quality, grid characteristics, and imaging characteristics, targeting can have issues. For instance, 1) We have found that large ice thickness gradients across individual atlas grid tiles can confuse Ptolemy’s per-tile normalization, causing square location predictions in seemingly random locations. 2) For grids where a majority of squares have cracks, Ptolemy may rank cracked squares higher than non-cracked squares. We hypothesize that the ML square ranking model becomes confounded because the brightness of a cracked square is greater than of a non-cracked square, yet still has other features of a non-cracked square, so it decides to weigh brightness highly. 3) For merged atlas tile images (Supplemental Figure 2), critical parameters such as square opening area may be distorted by any slight misalignment of the tiles, and the merged score, defined by the highest Ptolemy score, may not reflect the quality of the square if the image of the square had been intact in a tile.

The grouping in the sampling algorithm used in the experiments described here aimed to form groups with equal numbers of scored squares. We noticed that this grouping approach made it difficult to achieve the diversity of square areas required for *de novo* screening when the distribution of square areas itself is highly unbalanced. For example, if most squares on the grids are dry, the few squares with ice all end up in the same sampling group and would only be sampled once while dry squares would be sampled multiple times. To address this, we have added an option to use the area filter to place squares into a predefined area range. This eliminates the square area bias in sampling, but adds two extra user parameters that need to be defined by the grid mesh.

Using this new grouping, we have had some success in targeting squares on Spotiton^35, 36^ and chameleon^37^ nanowire grids which have characteristic stripes of sample across contiguous squares and no sample elsewhere on the grid. However, this approach is not reliable yet. We suspect that this is both because the percentage of such grids in the Ptolemy training set was low, and that usable squares do not usually have a significantly reduced area compared to the squares with no ice that make up the majority of squares. Work is in process to improve the success rate.

### Current limitations of the hole selection

In our experimental design for the timing and performance comparison between Autoscreen versus manual screening (Figure 2), no quality filtering was performed. In day-to-day operations, we have not found that these scores provided by Ptolemy are generally a better classifier than the ice thickness estimations. Additionally, the accuracy of the hole centers found by Ptolemy are compromised when holes are very different from the majority of the Ptolemy model training set. This includes cases when the Hole magnification images are taken at higher magnification so that the lattice is less clearly defined (e.g. when there are 4 or less holes in the image). The square lattice-based hole selection implemented in Ptolemy is also not able to target tilted or highly bent grids (Supplemental Figure 16), or lacey grids.

We are working on addressing these square and hole selection limitations so that Smart Leginon and Ptolemy generalize to as many grid and imaging characteristics as possible.

## Conclusion

We anticipate that Smart Leginon Autoscreen and associated functionality will significantly increase the throughput of cryoEM screening, an essential optimization step in the high-resolution single particle cryoEM pipeline. Simultaneously, the smart target selection algorithms significantly reduce operator time spent at the microscope, thus allowing for more time to be dedicated towards grid/sample preparation and data analysis. Moreover, idle microscope time outside of business hours (e.g. overnight and weekends) may be recovered, leading to an even greater effective increase in throughput. We envision that algorithms will continue to improve, particularly in the direction of real-time feedback from live-processing, which may enable intelligent, fully automated high-resolution collection that replicates or surpasses human performance.

## Acknowledgements

We thank Dr. Masahide Kikkawa (University of Tokyo) for the mApof sample and Dr. Brian Kloss (NYSBC) for expressing and purifying the mApof sample. We thank Andrew Santiago-Frangos, William Henriques, and Blake Wiedenheft (Montana State University) for allowing us to show their collection session in Figure 4, Goran Bajic (Icahn School of Medicine at Mount Sinai) for allowing us to show their collection session in Figure 5 and Supplemental Figure 15. We thank the SEMC IT team. Some of this work was performed at the Simons Electron Microscopy Center located at the New York Structural Biology Center, supported by grants from the Simons Foundation (SF349247) and NIH NIGMS (GM103310).

## Author Contributions Statement

A.C., P.K., C.S.P., B.C., T.B., and A.J.N. conceived of this project and designed the experiments, A.C., P.K., and T.B. designed and implemented the software, V.S. and K.N. prepared the sample, H.K., E.C., K.M., A.S., M.F.A., and K.N. tested the software, H.K., J.H.M., and E.C. analyzed the results, A.C., P.K., H.K., J.H.M., E.C., K.M., H.W., A.S., M.F.A., V.S., K.N., E.T.E., C.S.P., B.C., T.B., and A.J.N. analyzed results, and wrote and edited the manuscript.

## Software availability

Smart Leginon is freely and publicly available as two components: 1) Leginon and Autoscreen are in myami-3.6 release and above (http://leginon.org) and licensed under the Apache License, Version 2.0, and 2) Ptolemy is publicly available for academic use only (https://github.com/SMLC-NYSBC/ptolemy) and licensed under CC BY-NC 4.0. A tutorial for how to set up and use Smart Leginon and Autoscreen is available here: https://emg.nysbc.org/redmine/projects/leginon/wiki/Multi-grid_autoscreening

## Supplemental Information

**Supplemental Table 1:**
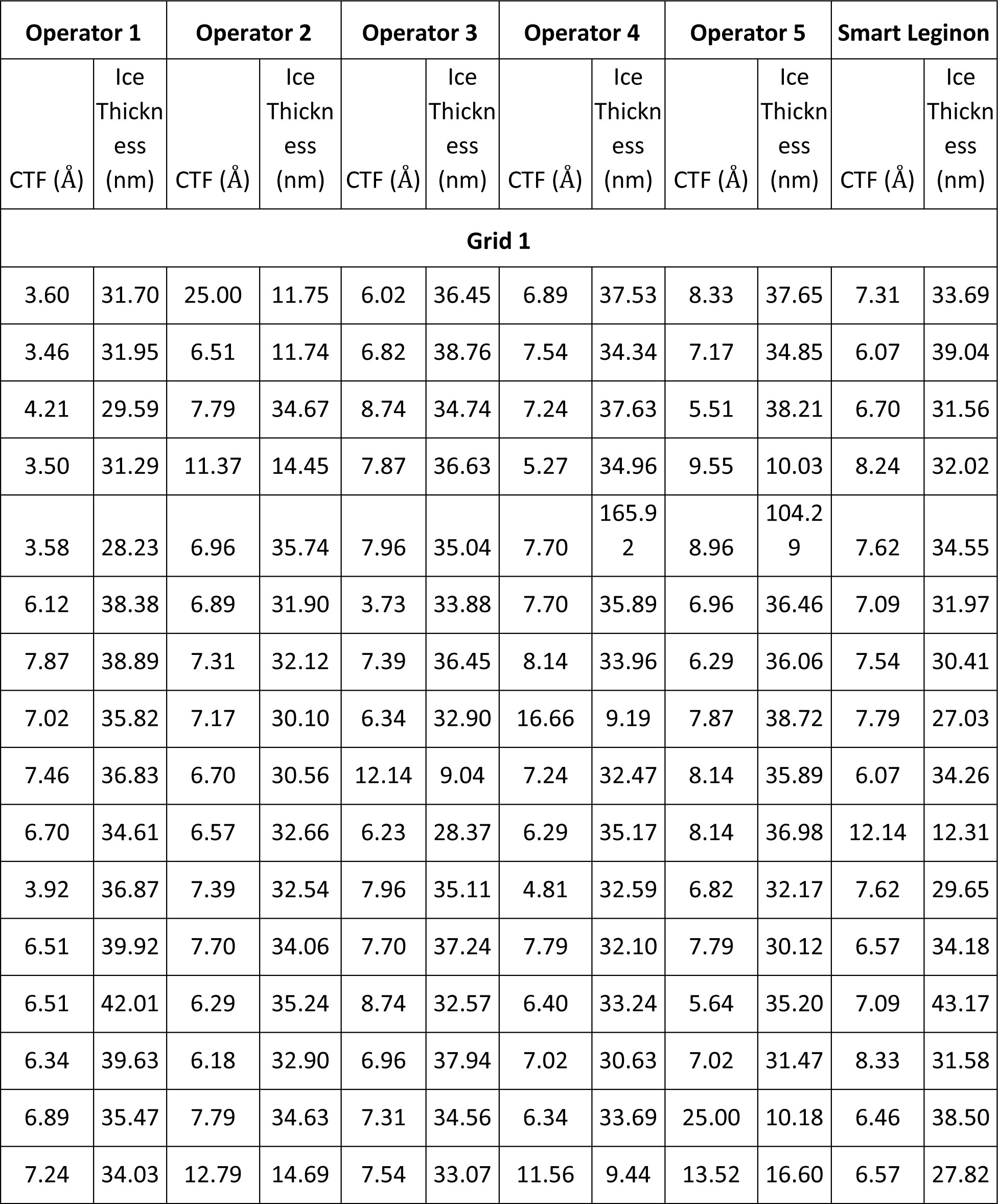
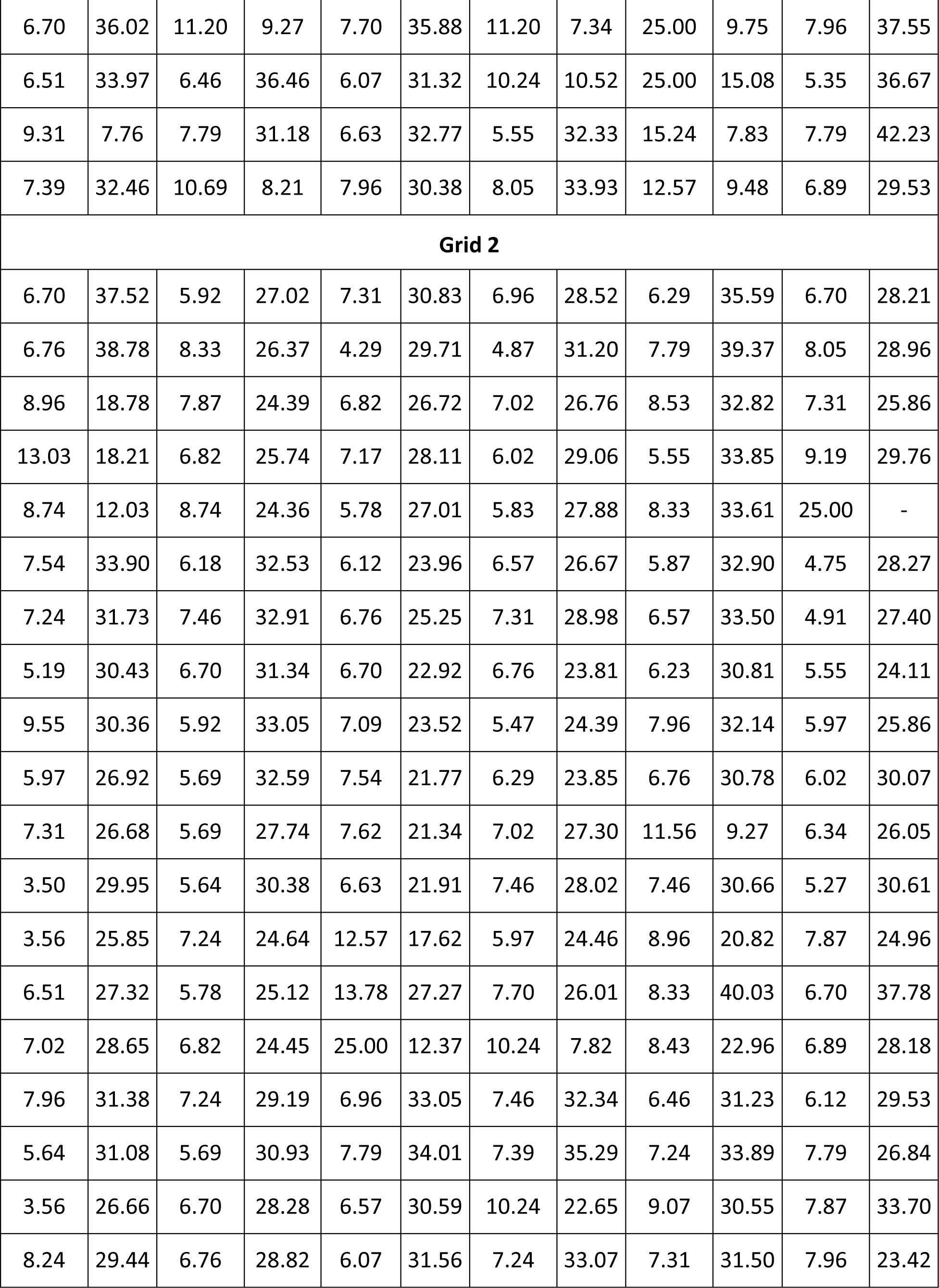
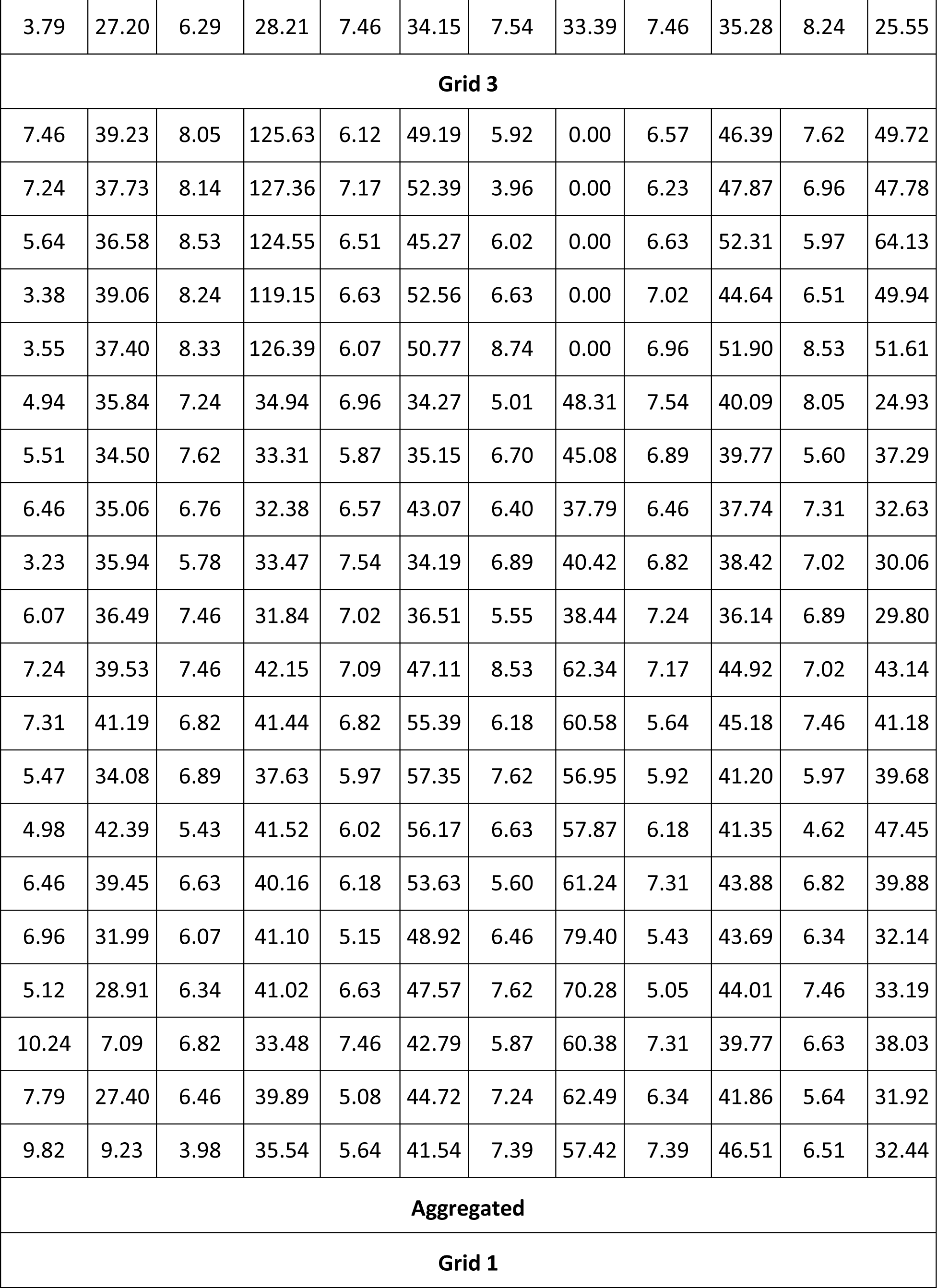
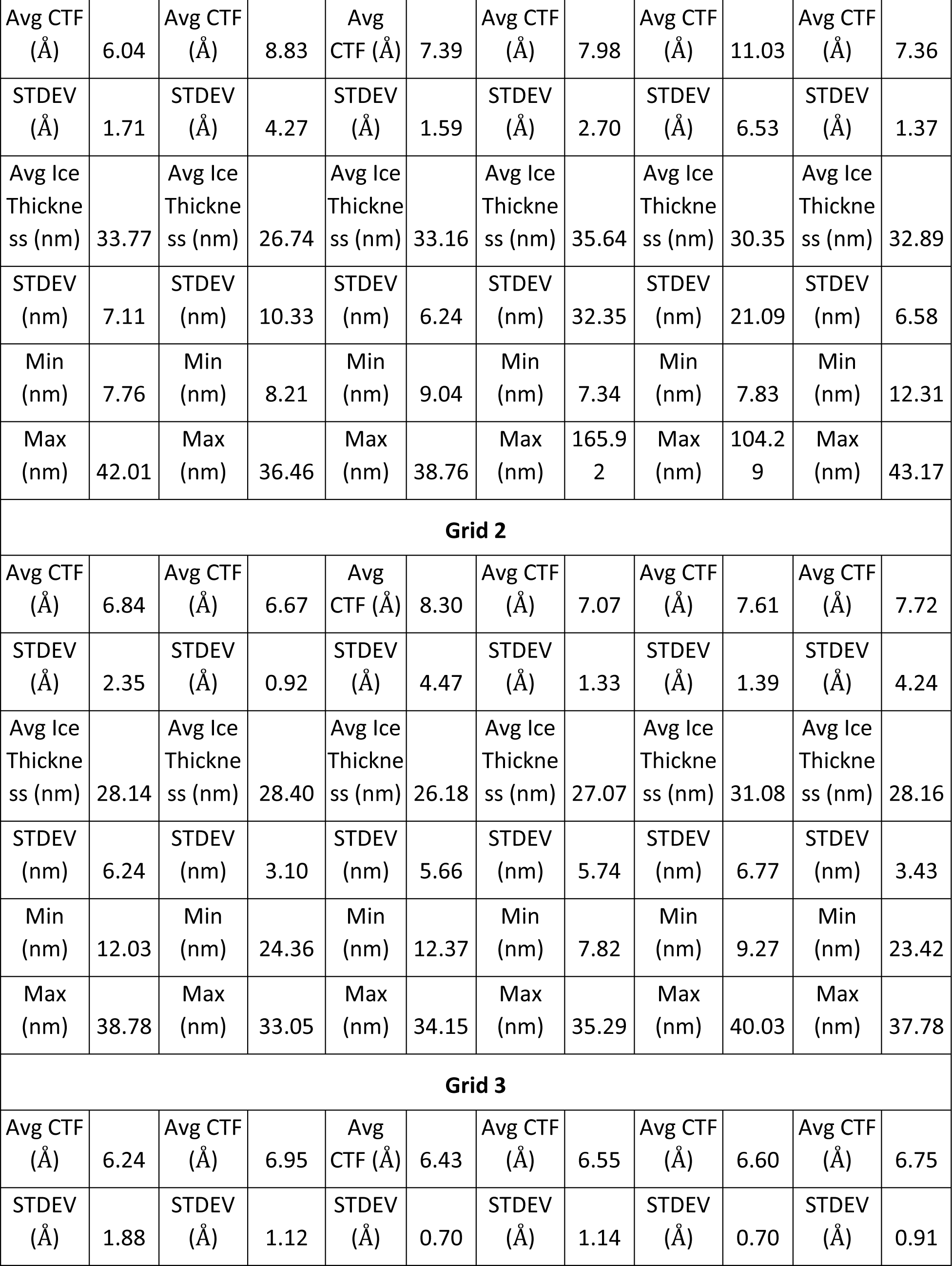
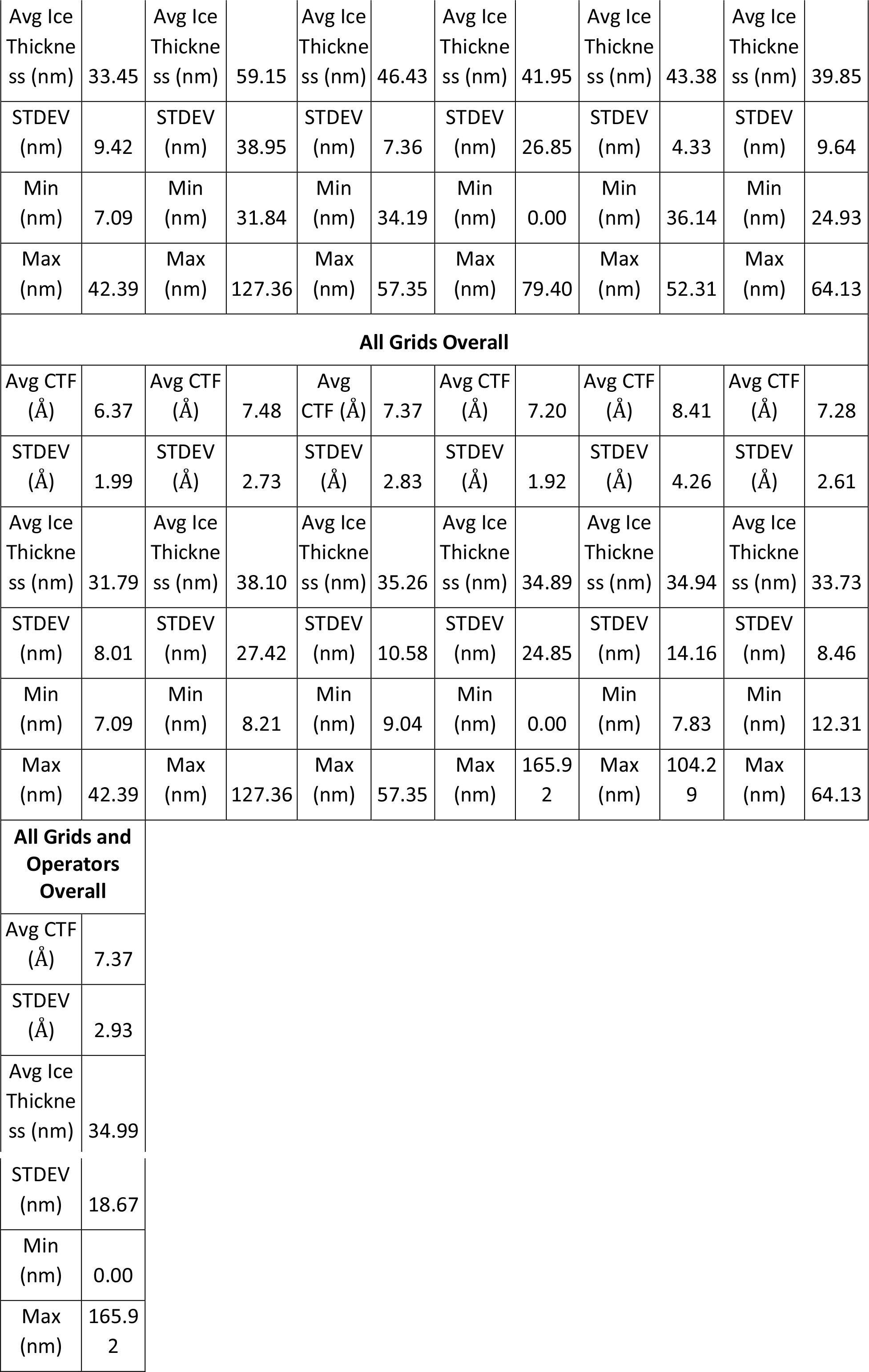
Exposure magnification CTF resolution and ice thickness comparisons. CTF resolution and ice thickness estimates along with averages and standard deviations for Exposure magnification images from the three mApof grids that were screened by Smart Leginon Autoscreen and the five microscope operators. Negative ice thickness estimates are reported as 0.00 nm. Note: Ice thickness calibration on the Glacios has an estimated error of about ±10 nm, so the values shown here should be considered as relative values, not absolute.

| Operator 1 |  | Operator 2 |  | Operator 3 |  | Operator 4 |  | Operator 5 |  | Smart Leginon |  |
| --- | --- | --- | --- | --- | --- | --- | --- | --- | --- | --- | --- |
| CTF (Å) | Ice Thickness (nm) | CTF (Å) | Ice Thickness (nm) | CTF (Å) | Ice Thickness (nm) | CTF (Å) | Ice Thickness (nm) | CTF (Å) | Ice Thickness (nm) | CTF (Å) | Ice Thickness (nm) |
| Grid 1 |  |  |  |  |  |  |  |  |  |  |  |
| 3.60 | 31.70 | 25.00 | 11.75 | 6.02 | 36.45 | 6.89 | 37.53 | 8.33 | 37.65 | 7.31 | 33.69 |
| 3.46 | 31.95 | 6.51 | 11.74 | 6.82 | 38.76 | 7.54 | 34.34 | 7.17 | 34.85 | 6.07 | 39.04 |
| 4.21 | 29.59 | 7.79 | 34.67 | 8.74 | 34.74 | 7.24 | 37.63 | 5.51 | 38.21 | 6.70 | 31.56 |
| 3.50 | 31.29 | 11.37 | 14.45 | 7.87 | 36.63 | 5.27 | 34.96 | 9.55 | 10.03 | 8.24 | 32.02 |
| 3.58 | 28.23 | 6.96 | 35.74 | 7.96 | 35.04 | 7.70 | 165.92 | 8.96 | 104.29 | 7.62 | 34.55 |
| 6.12 | 38.38 | 6.89 | 31.90 | 3.73 | 33.88 | 7.70 | 35.89 | 6.96 | 36.46 | 7.09 | 31.97 |
| 7.87 | 38.89 | 7.31 | 32.12 | 7.39 | 36.45 | 8.14 | 33.96 | 6.29 | 36.06 | 7.54 | 30.41 |
| 7.02 | 35.82 | 7.17 | 30.10 | 6.34 | 32.90 | 16.66 | 9.19 | 7.87 | 38.72 | 7.79 | 27.03 |
| 7.46 | 36.83 | 6.70 | 30.56 | 12.14 | 9.04 | 7.24 | 32.47 | 8.14 | 35.89 | 6.07 | 34.26 |
| 6.70 | 34.61 | 6.57 | 32.66 | 6.23 | 28.37 | 6.29 | 35.17 | 8.14 | 36.98 | 12.14 | 12.31 |
| 3.92 | 36.87 | 7.39 | 32.54 | 7.96 | 35.11 | 4.81 | 32.59 | 6.82 | 32.17 | 7.62 | 29.65 |
| 6.51 | 39.92 | 7.70 | 34.06 | 7.70 | 37.24 | 7.79 | 32.10 | 7.79 | 30.12 | 6.57 | 34.18 |
| 6.51 | 42.01 | 6.29 | 35.24 | 8.74 | 32.57 | 6.40 | 33.24 | 5.64 | 35.20 | 7.09 | 43.17 |
| 6.34 | 39.63 | 6.18 | 32.90 | 6.96 | 37.94 | 7.02 | 30.63 | 7.02 | 31.47 | 8.33 | 31.58 |
| 6.89 | 35.47 | 7.79 | 34.63 | 7.31 | 34.56 | 6.34 | 33.69 | 25.00 | 10.18 | 6.46 | 38.50 |
| 7.24 | 34.03 | 12.79 | 14.69 | 7.54 | 33.07 | 11.56 | 9.44 | 13.52 | 16.60 | 6.57 | 27.82 |
| 6.70 | 36.02 | 11.20 | 9.27 | 7.70 | 35.88 | 11.20 | 7.34 | 25.00 | 9.75 | 7.96 | 37.55 |
| 6.51 | 33.97 | 6.46 | 36.46 | 6.07 | 31.32 | 10.24 | 10.52 | 25.00 | 15.08 | 5.35 | 36.67 |
| 9.31 | 7.76 | 7.79 | 31.18 | 6.63 | 32.77 | 5.55 | 32.33 | 15.24 | 7.83 | 7.79 | 42.23 |
| 7.39 | 32.46 | 10.69 | 8.21 | 7.96 | 30.38 | 8.05 | 33.93 | 12.57 | 9.48 | 6.89 | 29.53 |
| **Grid 2** |  |  |  |  |  |  |  |  |  |  |  |
| 6.70 | 37.52 | 5.92 | 27.02 | 7.31 | 30.83 | 6.96 | 28.52 | 6.29 | 35.59 | 6.70 | 28.21 |
| 6.76 | 38.78 | 8.33 | 26.37 | 4.29 | 29.71 | 4.87 | 31.20 | 7.79 | 39.37 | 8.05 | 28.96 |
| 8.96 | 18.78 | 7.87 | 24.39 | 6.82 | 26.72 | 7.02 | 26.76 | 8.53 | 32.82 | 7.31 | 25.86 |
| 13.03 | 18.21 | 6.82 | 25.74 | 7.17 | 28.11 | 6.02 | 29.06 | 5.55 | 33.85 | 9.19 | 29.76 |
| 8.74 | 12.03 | 8.74 | 24.36 | 5.78 | 27.01 | 5.83 | 27.88 | 8.33 | 33.61 | 25.00 | - |
| 7.54 | 33.90 | 6.18 | 32.53 | 6.12 | 23.96 | 6.57 | 26.67 | 5.87 | 32.90 | 4.75 | 28.27 |
| 7.24 | 31.73 | 7.46 | 32.91 | 6.76 | 25.25 | 7.31 | 28.98 | 6.57 | 33.50 | 4.91 | 27.40 |
| 5.19 | 30.43 | 6.70 | 31.34 | 6.70 | 22.92 | 6.76 | 23.81 | 6.23 | 30.81 | 5.55 | 24.11 |
| 9.55 | 30.36 | 5.92 | 33.05 | 7.09 | 23.52 | 5.47 | 24.39 | 7.96 | 32.14 | 5.97 | 25.86 |
| 5.97 | 26.92 | 5.69 | 32.59 | 7.54 | 21.77 | 6.29 | 23.85 | 6.76 | 30.78 | 6.02 | 30.07 |
| 7.31 | 26.68 | 5.69 | 27.74 | 7.62 | 21.34 | 7.02 | 27.30 | 11.56 | 9.27 | 6.34 | 26.05 |
| 3.50 | 29.95 | 5.64 | 30.38 | 6.63 | 21.91 | 7.46 | 28.02 | 7.46 | 30.66 | 5.27 | 30.61 |
| 3.56 | 25.85 | 7.24 | 24.64 | 12.57 | 17.62 | 5.97 | 24.46 | 8.96 | 20.82 | 7.87 | 24.96 |
| 6.51 | 27.32 | 5.78 | 25.12 | 13.78 | 27.27 | 7.70 | 26.01 | 8.33 | 40.03 | 6.70 | 37.78 |
| 7.02 | 28.65 | 6.82 | 24.45 | 25.00 | 12.37 | 10.24 | 7.82 | 8.43 | 22.96 | 6.89 | 28.18 |
| 7.96 | 31.38 | 7.24 | 29.19 | 6.96 | 33.05 | 7.46 | 32.34 | 6.46 | 31.23 | 6.12 | 29.53 |
| 5.64 | 31.08 | 5.69 | 30.93 | 7.79 | 34.01 | 7.39 | 35.29 | 7.24 | 33.89 | 7.79 | 26.84 |
| 3.56 | 26.66 | 6.70 | 28.28 | 6.57 | 30.59 | 10.24 | 22.65 | 9.07 | 30.55 | 7.87 | 33.70 |
| 8.24 | 29.44 | 6.76 | 28.82 | 6.07 | 31.56 | 7.24 | 33.07 | 7.31 | 31.50 | 7.96 | 23.42 |
| 3.79 | 27.20 | 6.29 | 28.21 | 7.46 | 34.15 | 7.54 | 33.39 | 7.46 | 35.28 | 8.24 | 25.55 |
| **Grid 3** |  |  |  |  |  |  |  |  |  |  |  |
| 7.46 | 39.23 | 8.05 | 125.63 | 6.12 | 49.19 | 5.92 | 0.00 | 6.57 | 46.39 | 7.62 | 49.72 |
| 7.24 | 37.73 | 8.14 | 127.36 | 7.17 | 52.39 | 3.96 | 0.00 | 6.23 | 47.87 | 6.96 | 47.78 |
| 5.64 | 36.58 | 8.53 | 124.55 | 6.51 | 45.27 | 6.02 | 0.00 | 6.63 | 52.31 | 5.97 | 64.13 |
| 3.38 | 39.06 | 8.24 | 119.15 | 6.63 | 52.56 | 6.63 | 0.00 | 7.02 | 44.64 | 6.51 | 49.94 |
| 3.55 | 37.40 | 8.33 | 126.39 | 6.07 | 50.77 | 8.74 | 0.00 | 6.96 | 51.90 | 8.53 | 51.61 |
| 4.94 | 35.84 | 7.24 | 34.94 | 6.96 | 34.27 | 5.01 | 48.31 | 7.54 | 40.09 | 8.05 | 24.93 |
| 5.51 | 34.50 | 7.62 | 33.31 | 5.87 | 35.15 | 6.70 | 45.08 | 6.89 | 39.77 | 5.60 | 37.29 |
| 6.46 | 35.06 | 6.76 | 32.38 | 6.57 | 43.07 | 6.40 | 37.79 | 6.46 | 37.74 | 7.31 | 32.63 |
| 3.23 | 35.94 | 5.78 | 33.47 | 7.54 | 34.19 | 6.89 | 40.42 | 6.82 | 38.42 | 7.02 | 30.06 |
| 6.07 | 36.49 | 7.46 | 31.84 | 7.02 | 36.51 | 5.55 | 38.44 | 7.24 | 36.14 | 6.89 | 29.80 |
| 7.24 | 39.53 | 7.46 | 42.15 | 7.09 | 47.11 | 8.53 | 62.34 | 7.17 | 44.92 | 7.02 | 43.14 |
| 7.31 | 41.19 | 6.82 | 41.44 | 6.82 | 55.39 | 6.18 | 60.58 | 5.64 | 45.18 | 7.46 | 41.18 |
| 5.47 | 34.08 | 6.89 | 37.63 | 5.97 | 57.35 | 7.62 | 56.95 | 5.92 | 41.20 | 5.97 | 39.68 |
| 4.98 | 42.39 | 5.43 | 41.52 | 6.02 | 56.17 | 6.63 | 57.87 | 6.18 | 41.35 | 4.62 | 47.45 |
| 6.46 | 39.45 | 6.63 | 40.16 | 6.18 | 53.63 | 5.60 | 61.24 | 7.31 | 43.88 | 6.82 | 39.88 |
| 6.96 | 31.99 | 6.07 | 41.10 | 5.15 | 48.92 | 6.46 | 79.40 | 5.43 | 43.69 | 6.34 | 32.14 |
| 5.12 | 28.91 | 6.34 | 41.02 | 6.63 | 47.57 | 7.62 | 70.28 | 5.05 | 44.01 | 7.46 | 33.19 |
| 10.24 | 7.09 | 6.82 | 33.48 | 7.46 | 42.79 | 5.87 | 60.38 | 7.31 | 39.77 | 6.63 | 38.03 |
| 7.79 | 27.40 | 6.46 | 39.89 | 5.08 | 44.72 | 7.24 | 62.49 | 6.34 | 41.86 | 5.64 | 31.92 |
| 9.82 | 9.23 | 3.98 | 35.54 | 5.64 | 41.54 | 7.39 | 57.42 | 7.39 | 46.51 | 6.51 | 32.44 |
| **Aggregated** |  |  |  |  |  |  |  |  |  |  |  |
| **Grid 1** |  |  |  |  |  |  |  |  |  |  |  |
| Avg CTF (Å) | 6.04 | Avg CTF (Å) | 8.83 | Avg CTF (Å) | 7.39 | Avg CTF (Å) | 7.98 | Avg CTF (Å) | 11.03 | Avg CTF (Å) | 7.36 |
| STDEV (Å) | 1.71 | STDEV (Å) | 4.27 | STDEV (Å) | 1.59 | STDEV (Å) | 2.70 | STDEV (Å) | 6.53 | STDEV (Å) | 1.37 |
| Avg Ice Thickne ss (nm) | 33.77 | Avg Ice Thickne ss (nm) | 26.74 | Avg Ice Thickne ss (nm) | 33.16 | Avg Ice Thickne ss (nm) | 35.64 | Avg Ice Thickne ss (nm) | 30.35 | Avg Ice Thickne ss (nm) | 32.89 |
| STDEV (nm) | 7.11 | STDEV (nm) | 10.33 | STDEV (nm) | 6.24 | STDEV (nm) | 32.35 | STDEV (nm) | 21.09 | STDEV (nm) | 6.58 |
| Min (nm) | 7.76 | Min (nm) | 8.21 | Min (nm) | 9.04 | Min (nm) | 7.34 | Min (nm) | 7.83 | Min (nm) | 12.31 |
| Max (nm) | 42.01 | Max (nm) | 36.46 | Max (nm) | 38.76 | Max (nm) | 165.92 | Max (nm) | 104.29 | Max (nm) | 43.17 |
| **Grid 2** |  |  |  |  |  |  |  |  |  |  |  |
| Avg CTF (Å) | 6.84 | Avg CTF (Å) | 6.67 | Avg CTF (Å) | 8.30 | Avg CTF (Å) | 7.07 | Avg CTF (Å) | 7.61 | Avg CTF (Å) | 7.72 |
| STDEV (Å) | 2.35 | STDEV (Å) | 0.92 | STDEV (Å) | 4.47 | STDEV (Å) | 1.33 | STDEV (Å) | 1.39 | STDEV (Å) | 4.24 |
| Avg Ice Thickne ss (nm) | 28.14 | Avg Ice Thickne ss (nm) | 28.40 | Avg Ice Thickne ss (nm) | 26.18 | Avg Ice Thickne ss (nm) | 27.07 | Avg Ice Thickne ss (nm) | 31.08 | Avg Ice Thickne ss (nm) | 28.16 |
| STDEV (nm) | 6.24 | STDEV (nm) | 3.10 | STDEV (nm) | 5.66 | STDEV (nm) | 5.74 | STDEV (nm) | 6.77 | STDEV (nm) | 3.43 |
| Min (nm) | 12.03 | Min (nm) | 24.36 | Min (nm) | 12.37 | Min (nm) | 7.82 | Min (nm) | 9.27 | Min (nm) | 23.42 |
| Max (nm) | 38.78 | Max (nm) | 33.05 | Max (nm) | 34.15 | Max (nm) | 35.29 | Max (nm) | 40.03 | Max (nm) | 37.78 |
| **Grid 3** |  |  |  |  |  |  |  |  |  |  |  |
| Avg CTF (Å) | 6.24 | Avg CTF (Å) | 6.95 | Avg CTF (Å) | 6.43 | Avg CTF (Å) | 6.55 | Avg CTF (Å) | 6.60 | Avg CTF (Å) | 6.75 |
| STDEV (Å) | 1.88 | STDEV (Å) | 1.12 | STDEV (Å) | 0.70 | STDEV (Å) | 1.14 | STDEV (Å) | 0.70 | STDEV (Å) | 0.91 |
| Avg Ice Thickness (nm) | 33.45 | Avg Ice Thickness (nm) | 59.15 | Avg Ice Thickness (nm) | 46.43 | Avg Ice Thickness (nm) | 41.95 | Avg Ice Thickness (nm) | 43.38 | Avg Ice Thickness (nm) | 39.85 |
| STDEV (nm) | 9.42 | STDEV (nm) | 38.95 | STDEV (nm) | 7.36 | STDEV (nm) | 26.85 | STDEV (nm) | 4.33 | STDEV (nm) | 9.64 |
| Min (nm) | 7.09 | Min (nm) | 31.84 | Min (nm) | 34.19 | Min (nm) | 0.00 | Min (nm) | 36.14 | Min (nm) | 24.93 |
| Max (nm) | 42.39 | Max (nm) | 127.36 | Max (nm) | 57.35 | Max (nm) | 79.40 | Max (nm) | 52.31 | Max (nm) | 64.13 |
| **All Grids Overall** |  |  |  |  |  |  |  |  |  |  |  |
| Avg CTF (Å) | 6.37 | Avg CTF (Å) | 7.48 | Avg CTF (Å) | 7.37 | Avg CTF (Å) | 7.20 | Avg CTF (Å) | 8.41 | Avg CTF (Å) | 7.28 |
| STDEV (Å) | 1.99 | STDEV (Å) | 2.73 | STDEV (Å) | 2.83 | STDEV (Å) | 1.92 | STDEV (Å) | 4.26 | STDEV (Å) | 2.61 |
| Avg Ice Thickness (nm) | 31.79 | Avg Ice Thickness (nm) | 38.10 | Avg Ice Thickness (nm) | 35.26 | Avg Ice Thickness (nm) | 34.89 | Avg Ice Thickness (nm) | 34.94 | Avg Ice Thickness (nm) | 33.73 |
| STDEV (nm) | 8.01 | STDEV (nm) | 27.42 | STDEV (nm) | 10.58 | STDEV (nm) | 24.85 | STDEV (nm) | 14.16 | STDEV (nm) | 8.46 |
| Min (nm) | 7.09 | Min (nm) | 8.21 | Min (nm) | 9.04 | Min (nm) | 0.00 | Min (nm) | 7.83 | Min (nm) | 12.31 |
| Max (nm) | 42.39 | Max (nm) | 127.36 | Max (nm) | 57.35 | Max (nm) | 165.92 | Max (nm) | 104.29 | Max (nm) | 64.13 |
| **All Grids and Operators Overall** |  |  |  |  |  |  |  |  |  |  |  |
| Avg CTF (Å) | 7.37 |  |  |  |  |  |  |  |  |  |  |
| STDEV (Å) | 2.93 |  |  |  |  |  |  |  |  |  |  |
| Avg Ice Thickness (nm) | 34.99 |  |  |  |  |  |  |  |  |  |  |

**Supplemental Table 1 |** CTF resolution and ice thickness estimates along with averages and standard deviations for Exposure magnification images from the three mApof grids that were screened by Smart Leginon Autoscreen and the five microscope operators.
|  |  |
| --- | --- |
| STDEV<br>(nm) | 18.67 |
| Min<br>(nm) | 0.00 |
| Max<br>(nm) | 165.9<br>2 |

**Supplemental Figure 1:**
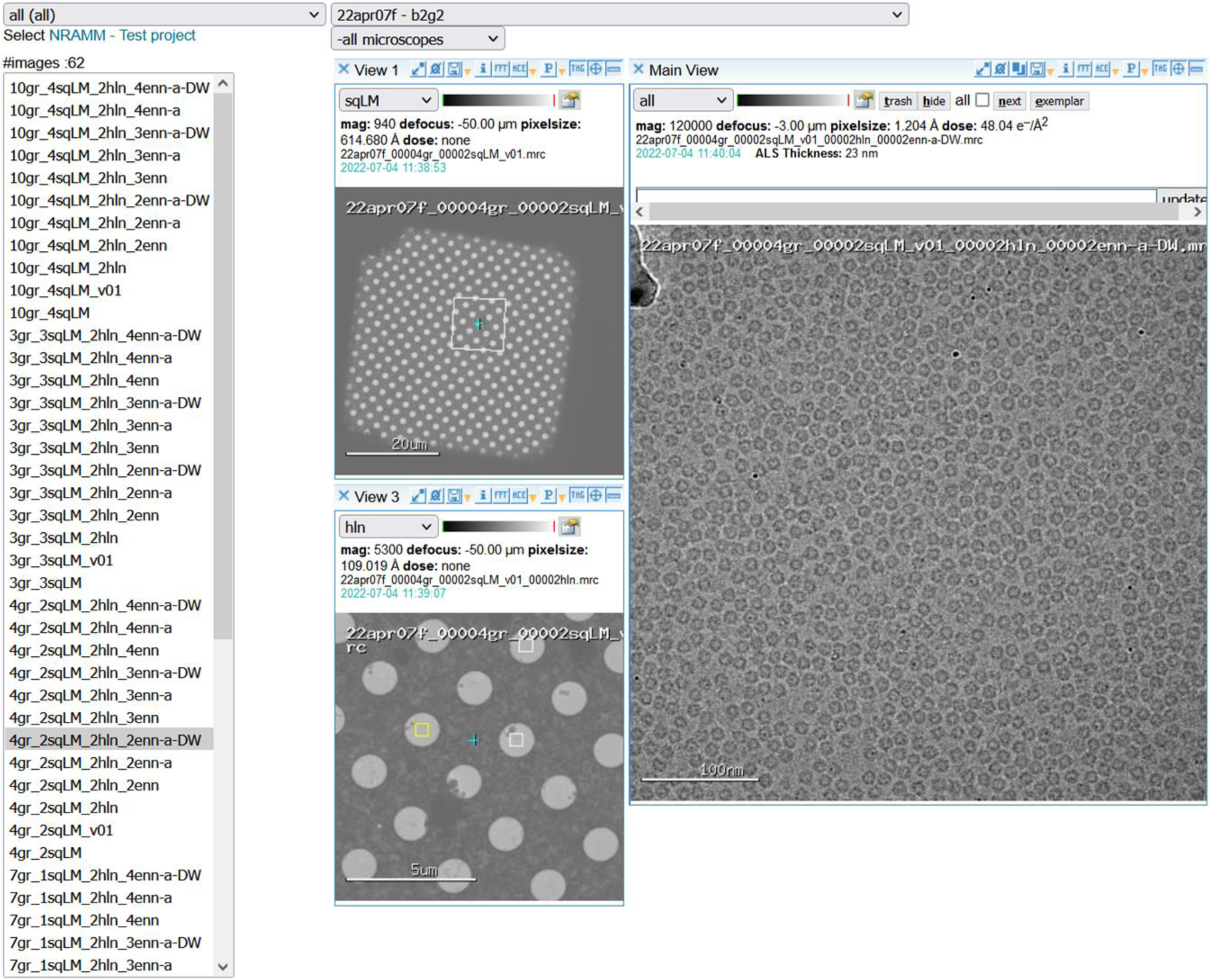
Multi-scale imaging (MSI) visualization with the Appion 3 Way Viewer. The Appion 3 Way Viewer allows for rapid visualization of a full set of MSI images; the atlas (not shown), the Square magnification image (top-left), the corresponding Hole magnification image (bottom-left), and the corresponding Exposure magnification image (right).

**Supplemental Figure 2:**
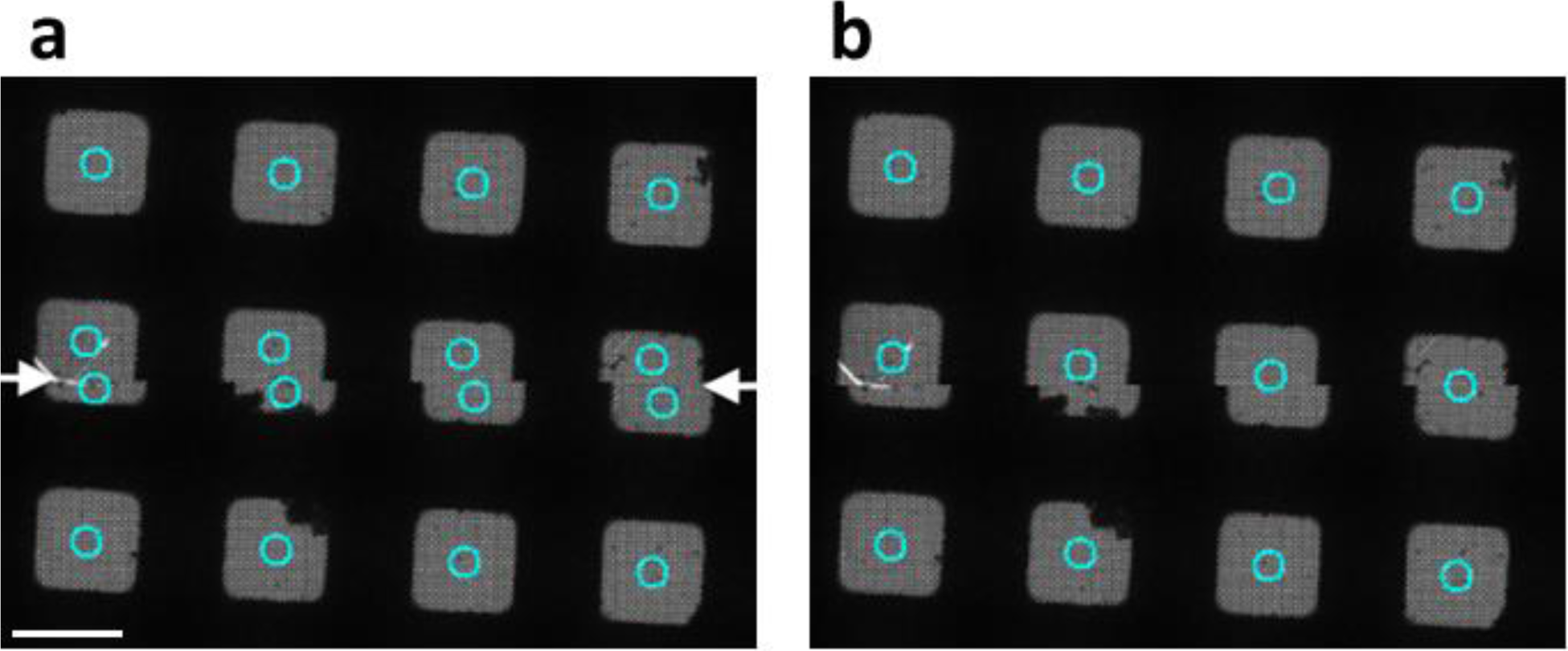
Smart Leginon atlas grid tile image merging. The atlas grid tile image merging update to Leginon. **(a)** Two tiles - top and bottom halves of the image - where Ptolemy returned square locations and metadata for each image. The squares at the edge of the tiles end up with two sets of square locations and metadata. **(b)** The tiles after square location and metadata merging. Scale bar is 50 μm.

**Supplemental Figure 3:**
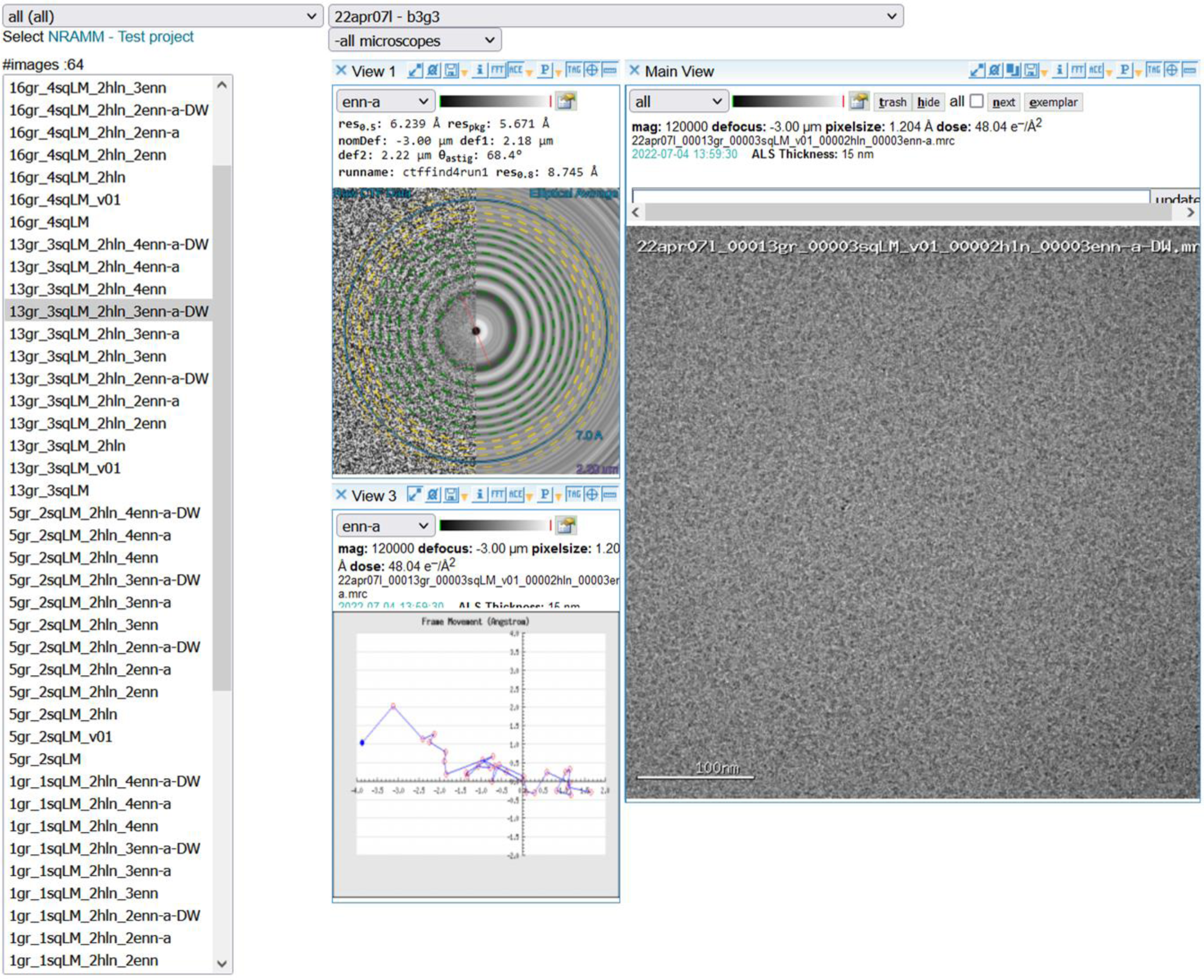
Appion 3 Way Viewer for analyzing pre-processing results. The Appion 3 Way Viewer allows for near real-time analysis of Exposure magnification micrograph quality, including ice thickness estimation, CTF estimation, and frame alignment results, as shown here.

**Supplemental Figure 4:**
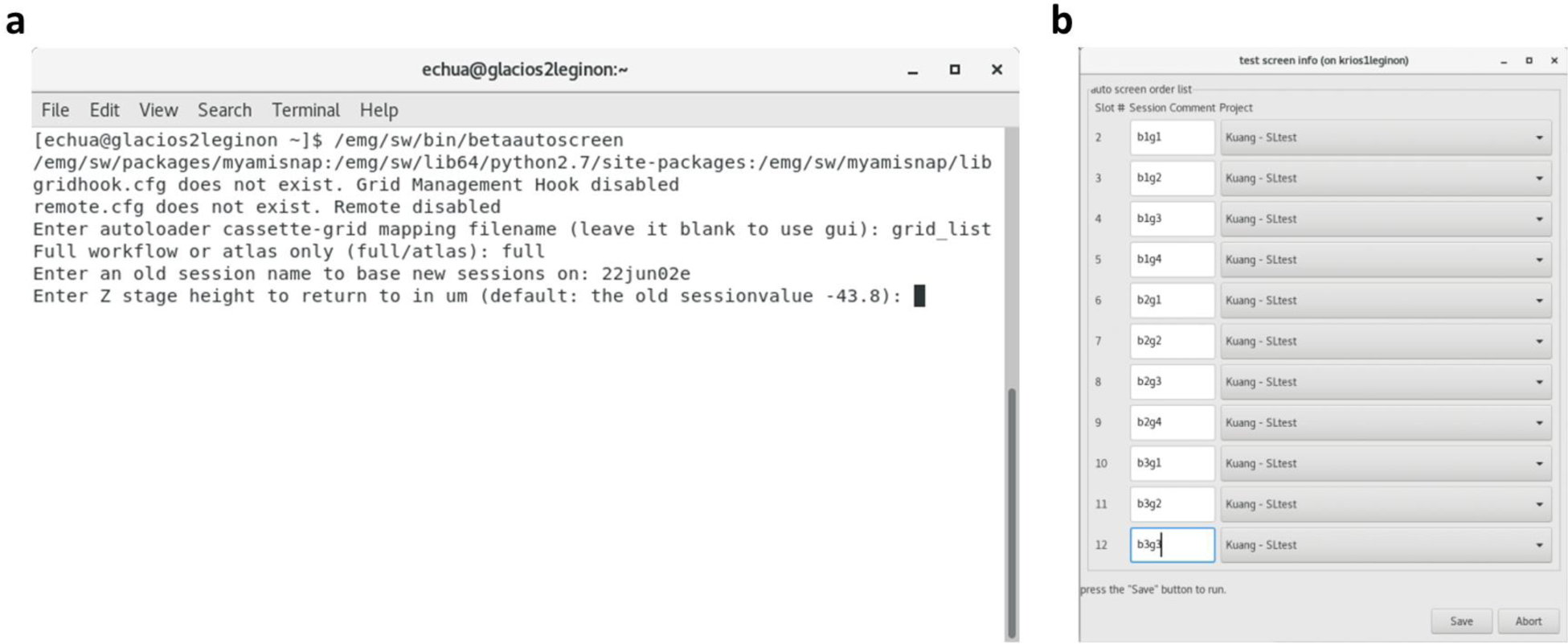
Smart Leginon Autoscreen command line workflow. Smart Leginon Autoscreen is executed by command line where the operator is asked to enter four pieces of information **(a)**, then Autoscreen will screen all grids in the microscope completely unattended. The first piece of information is the names of each grid and corresponding projects that Leginon will associate them with; names can be listed in a file and the filename can be entered as in **(a)**, or the entry can be left blank to open a GUI **(b)**. Next, the operator selects whether to perform the full MSI workflow on all grids, or to only collect atlases of all grids. Next, the operator inputs the name of an example Smart Leginon session from which all settings will be imported for screening all grids. Lastly, the default stage Z height may be changed, if necessary.

**Supplemental Figure 5:**
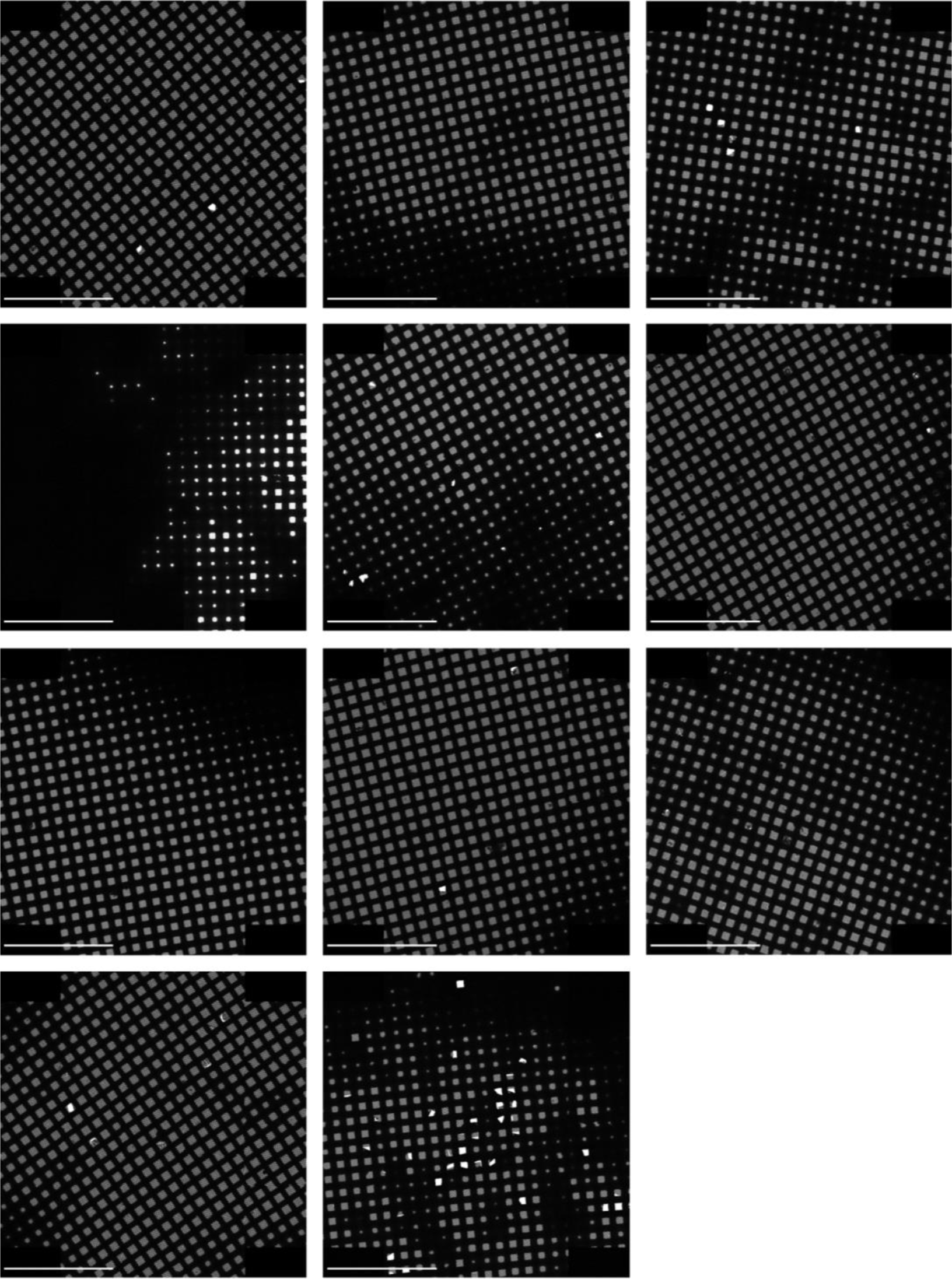
Atlases collected by Smart Leginon Autoscreen. Atlas images of 11 grids after Smart Leginon Autoscreen allowed for the rapid identification of 8 good to excellent grids (made by one person) and 3 poor grids (made by another). Scale bars are 500 μm.

**Supplemental Figure 6:**
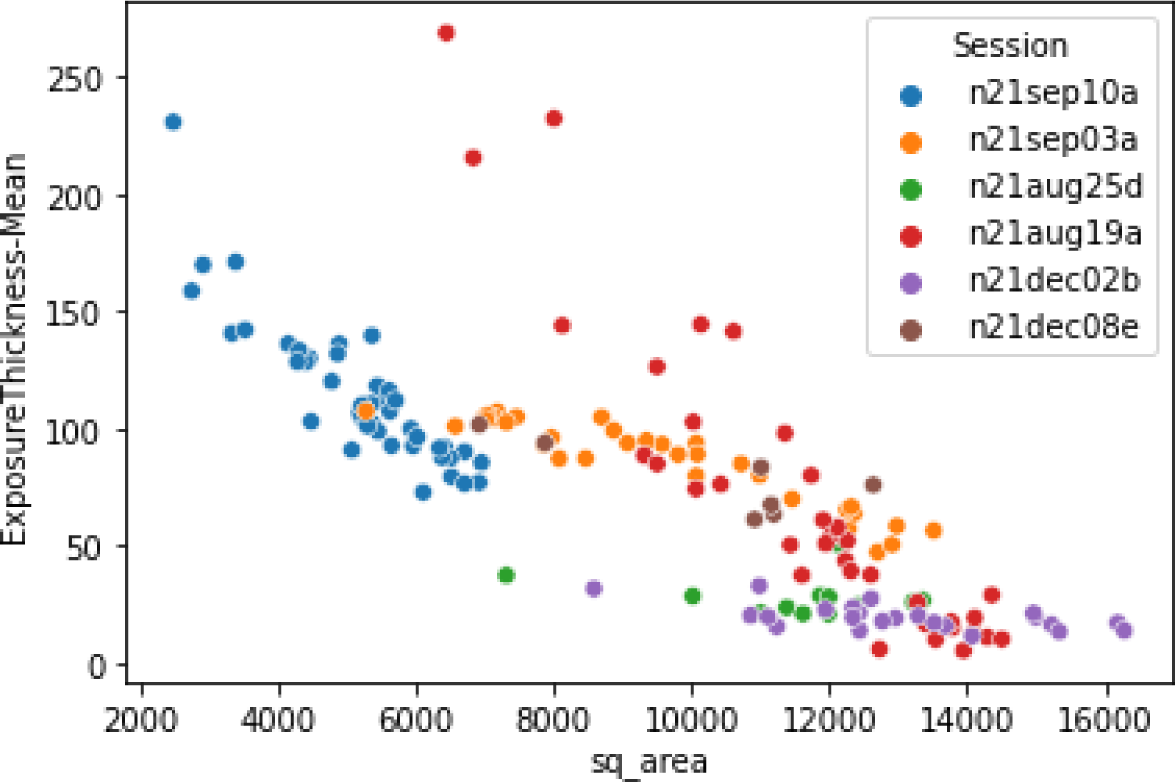
Square area correlates with ice thickness. Average ice thickness (nm) of holes in squares estimated from Exposure magnification images using an energy filter for calibration (y-axis) versus square area (arbitrary units) as calculated by Ptolemy (x-axis) across several collection sessions of different grids. In general, the larger the square area, the thinner the ice.

**Supplemental Figure 7:**
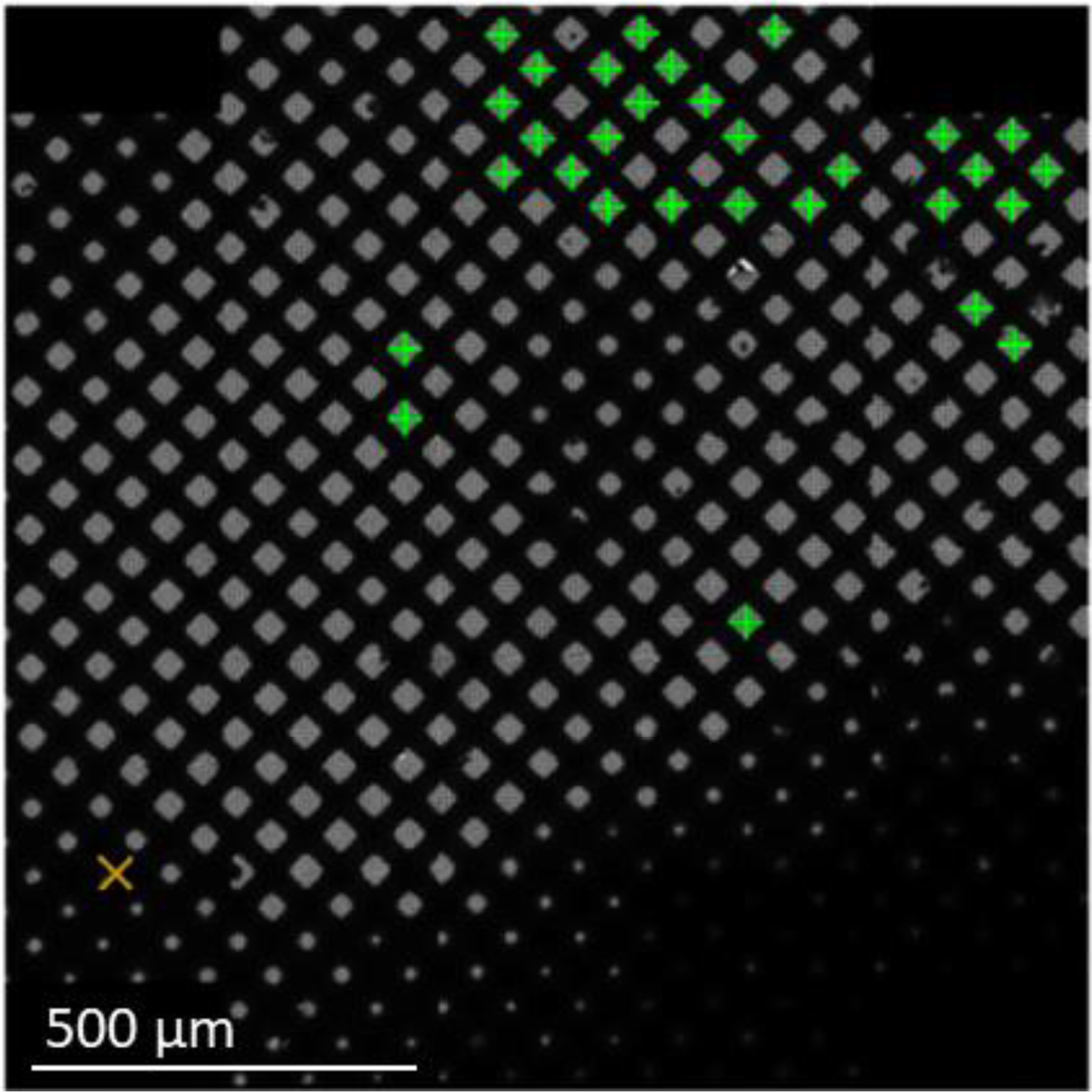
Smart Leginon can screen squares of a particular area. Smart Leginon may be used to screen squares in a particular area range. In this example, the user has prior knowledge that their particles behave well in large squares which correspond to thin ice, thus Smart Leginon was set up to screen the 30 highest-ranked large squares (green plus symbols, +).

**Supplemental Figure 8:**
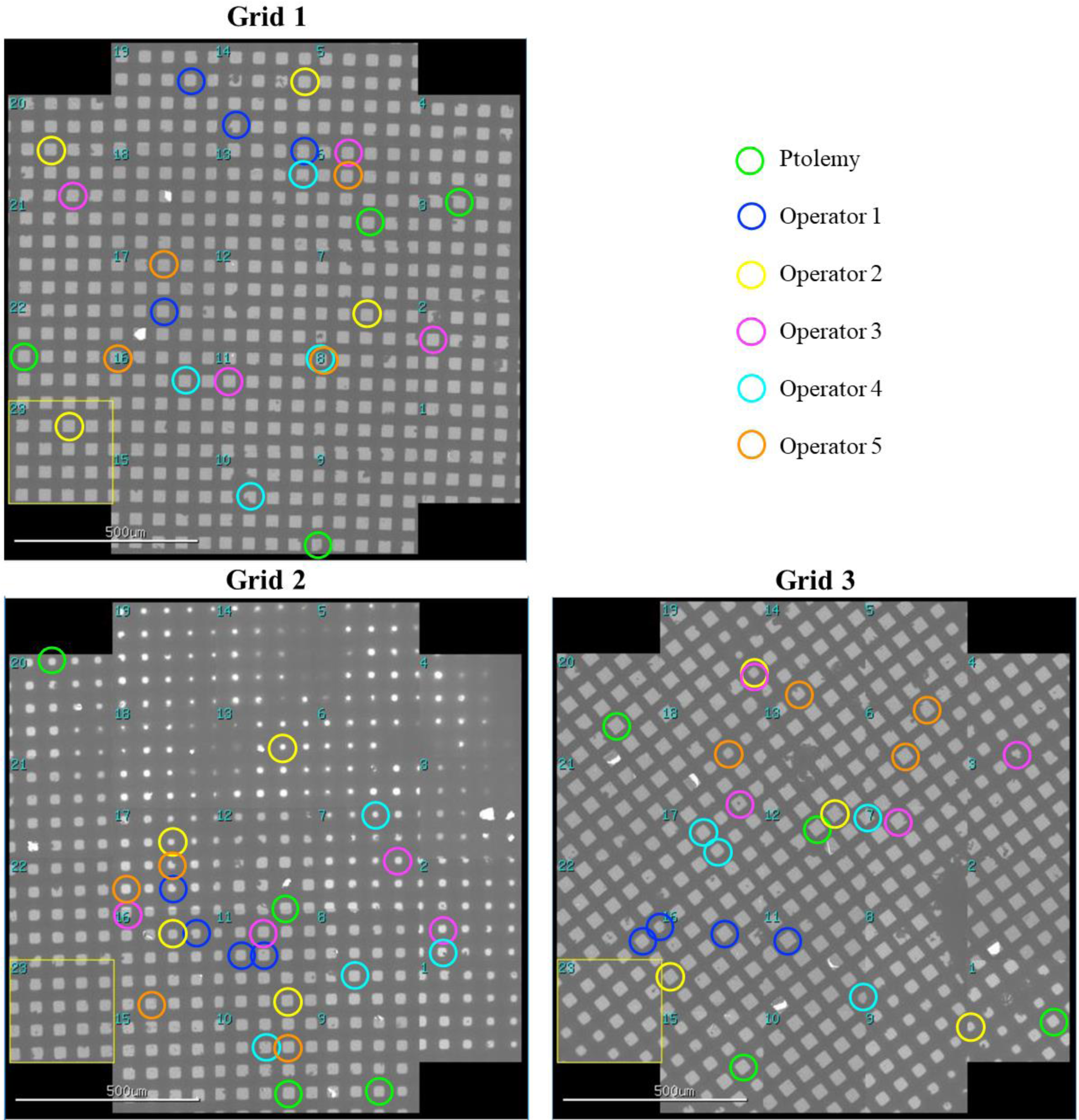
Comparison between Smart Leginon Autoscreen and Operators’ independent square targeting. A comparison of three unseen grids between Smart Leginon targeting and five expert microscope operators’ independent targeting. For each grid, Smart Leginon and the operators were instructed to choose the ‘best’ square from four equal-sized groups of square areas. Scale bars are 500 μm.

**Supplemental Figure 9:**
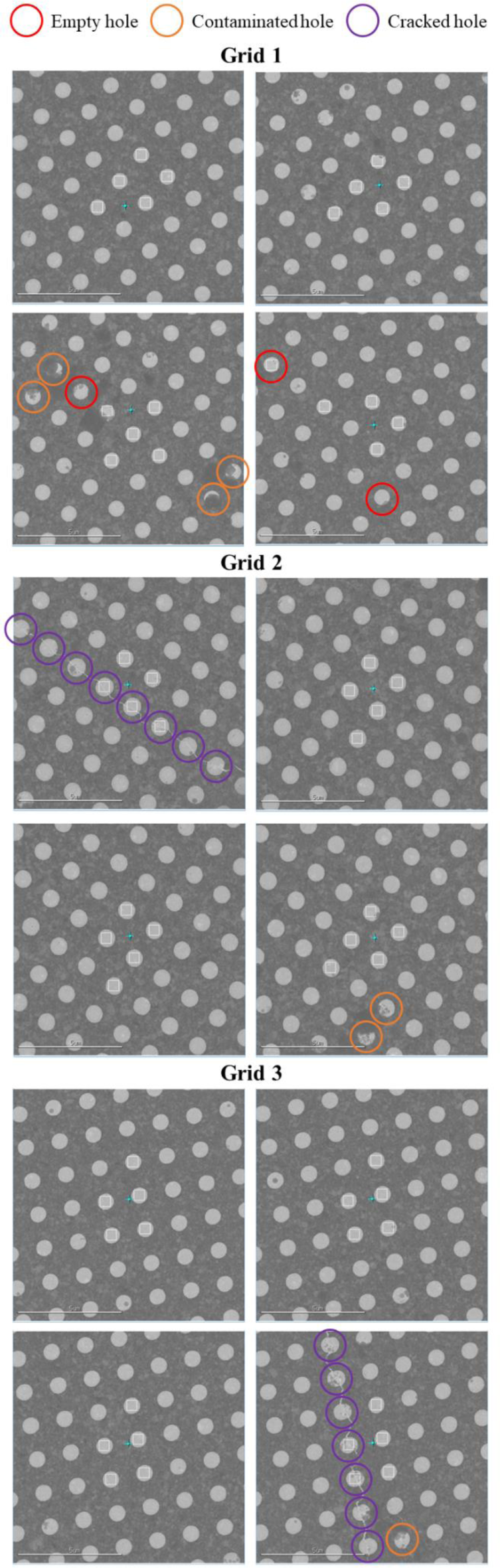
Analysis of holes from each square image for Operator 1. Empty, contaminated, and cracked holes are identified for four squares from three grids collected by operator 1. The remaining holes are considered ‘good’. Scale bars are 5 μm.

**Supplemental Figure 10:**
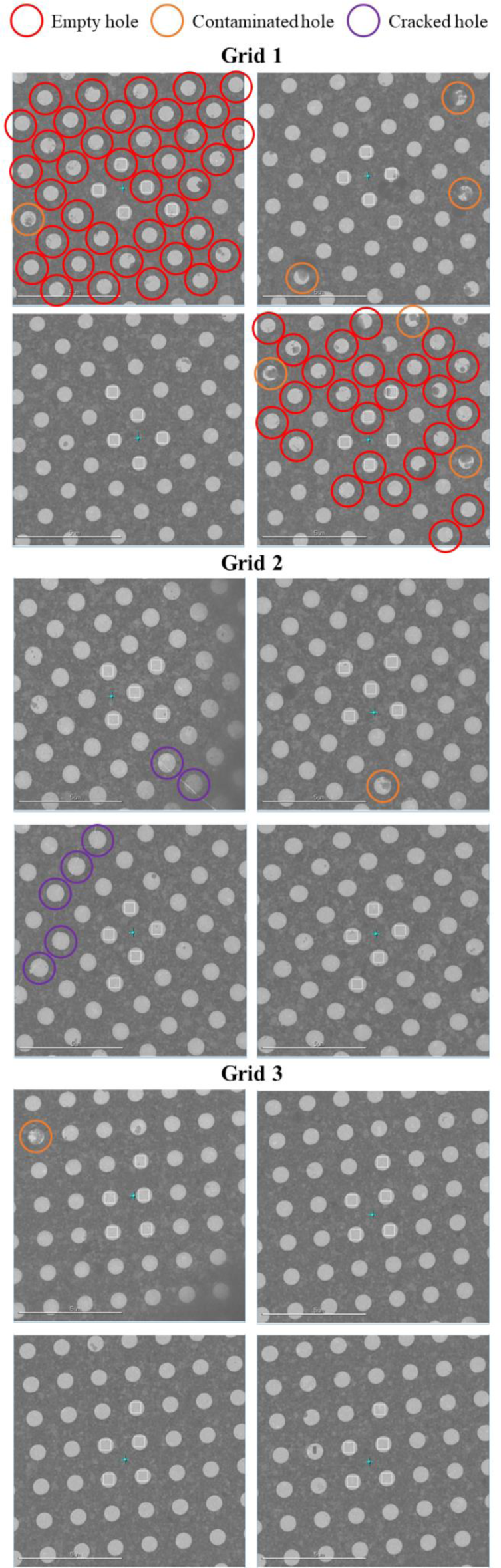
Analysis of holes from each square image for Operator 2. Empty, contaminated, and cracked holes are identified for four squares from three grids collected by operator 2. The remaining holes are considered ‘good’. Scale bars are 5 μm.

**Supplemental Figure 11:**
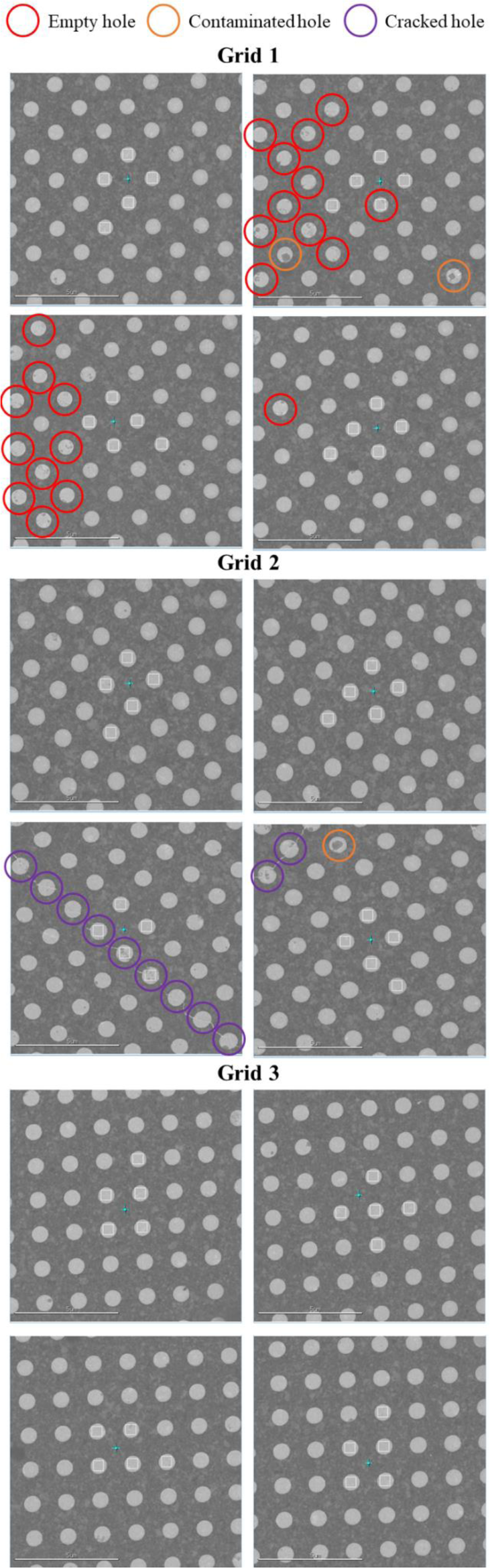
Analysis of holes from each square image for Operator 3. Empty, contaminated, and cracked holes are identified for four squares from three grids collected by operator 3. The remaining holes are considered ‘good’. Scale bars are 5 μm.

**Supplemental Figure 12:**
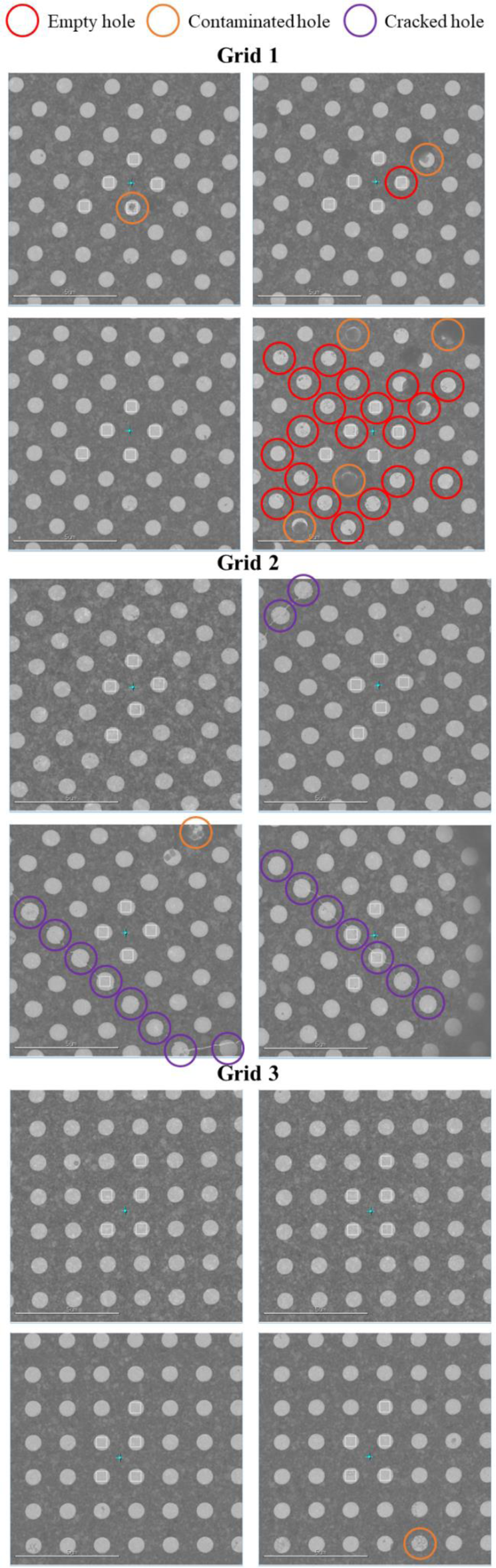
Analysis of holes from each square image for Operator 4. Empty, contaminated, and cracked holes are identified for four squares from three grids collected by operator 4. The remaining holes are considered ‘good’. Scale bars are 5 μm.

**Supplemental Figure 13:**
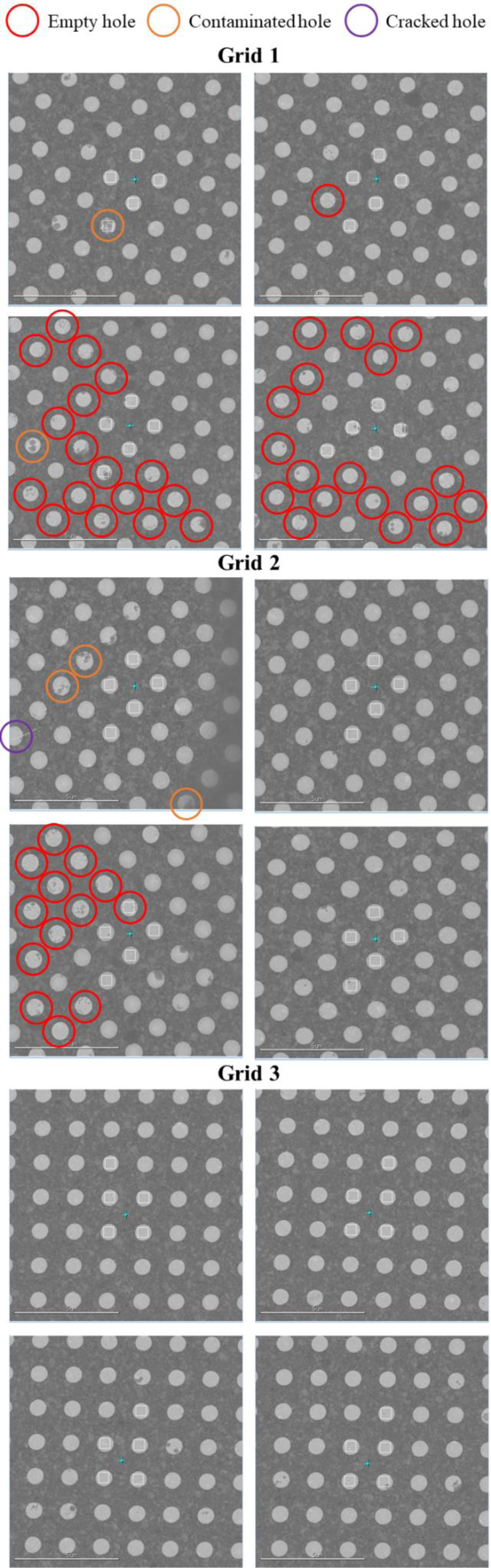
Analysis of holes from each square image for Operator 5. Empty, contaminated, and cracked holes are identified for four squares from three grids collected by operator 5. The remaining holes are considered ‘good’. Scale bars are 5 μm.

**Supplemental Figure 14:**
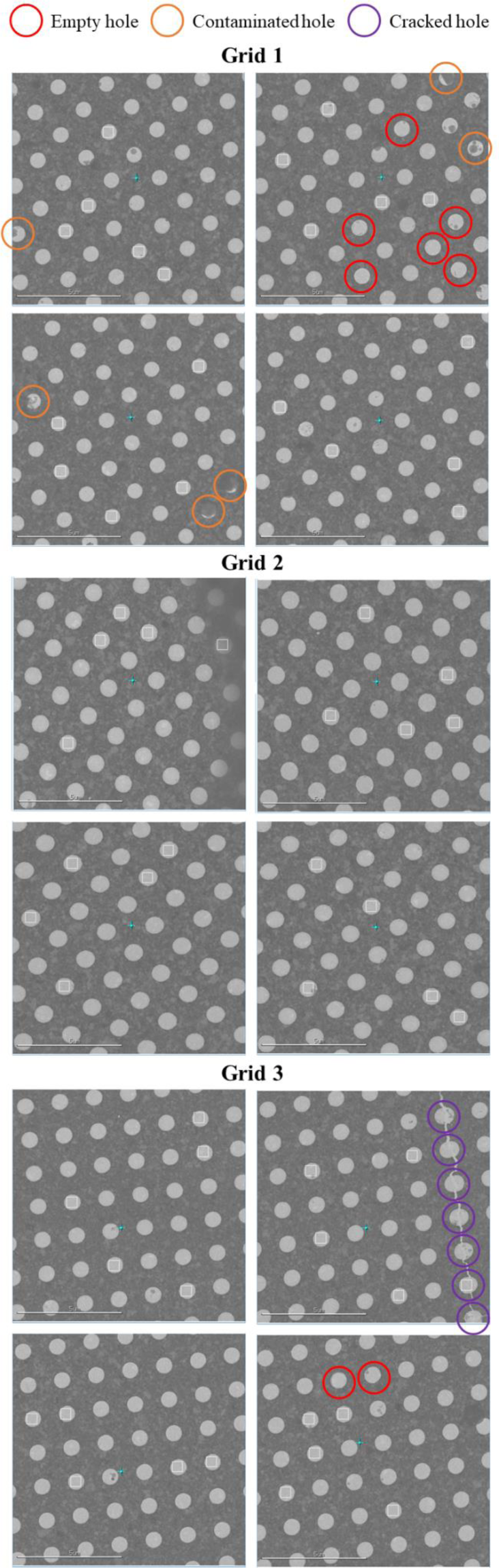
Analysis of holes from each square image for Smart Leginon Autoscreen. Empty, contaminated, and cracked holes are identified for four squares from three grids collected by Smart Leginon Autoscreen. The remaining holes are considered ‘good’. Scale bars are 5 μm.

**Supplemental Figure 15:**
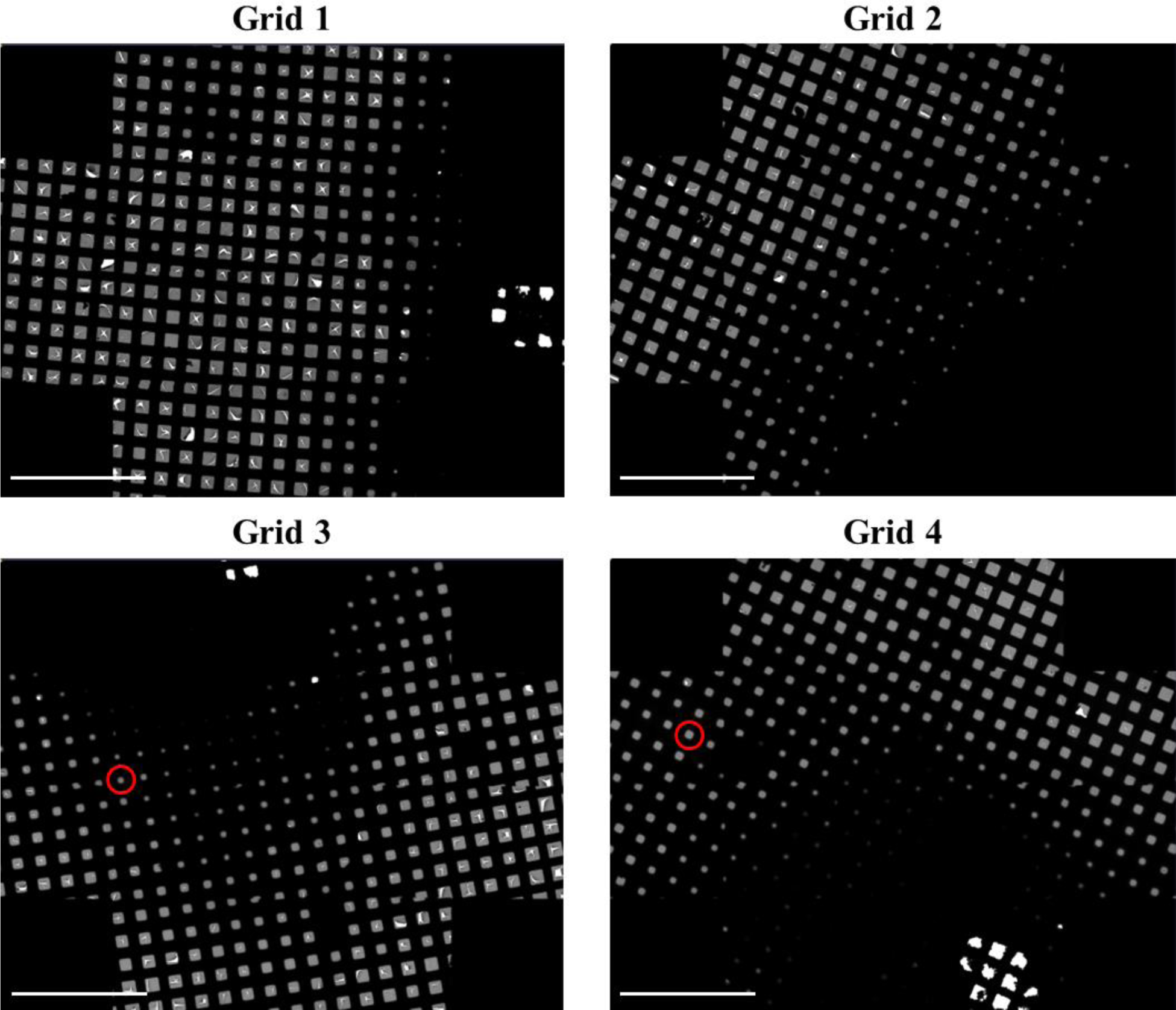
Four user grids screened on a Krios with Smart Leginon before collection. Four user grids, ordered by the user from best to worst, were screened with Smart Leginon prior to setting up a 43-hour high-resolution collection. The 2-hour Smart Leginon MSI screening analysis determined that in fact the proper order from best to worst was Grid 4, Grid 3, then either Grid 2 or Grid 1. The squares circled in red on Grid 3 & 4 show example squares that were found to be optimal for collection. Several squares with different areas (Figure 5) needed to be screened on each grid before making this determination. Scale bars are 500 μm

**Supplemental Figure 16:**
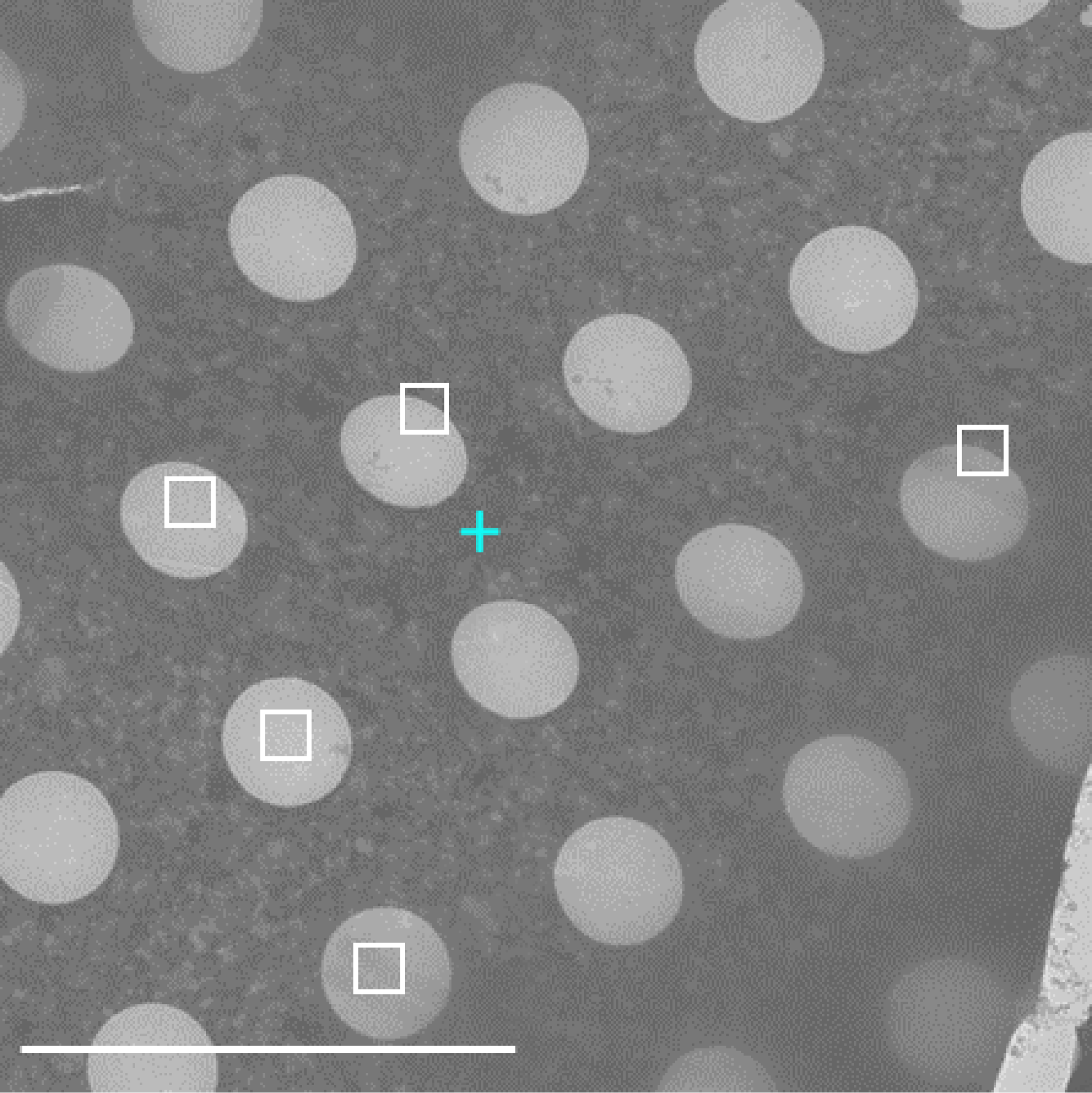
Smart Leginon failure case at hole level. A current limitation in Smart Leginon hole identification. Ptolemy hole lattice targeting on a grid with bent gold substrate shows that Ptolemy does not target well for non-square lattices. Scale bar is 5 μm.

